# Towards a scaled-up T cell-mediated cytotoxicity assay in 3D cell culture using microscopy

**DOI:** 10.1101/842039

**Authors:** Elinor Gottschalk, Eric Czech, Bulent Arman Aksoy, Pinar Aksoy, Jeff Hammerbacher

## Abstract

Three-dimensional (3D) cell culture systems with tumor spheroids are being adopted for research on the antitumor activity of drug treatments and cytotoxic T cells. Analysis of the cytotoxic effect on 3D tumor cultures within a 3D scaffold, such as collagen, is challenging. Image-based approaches often use confocal microscopy, which greatly limits the sample size of tumor spheroids that can be assayed. We explored a system where tumor spheroids growing in a collagen gel within a microfluidics chip can be treated with drugs or co-cultured with T cells. We attempted to adapt the system to measure the death of cells in the tumor spheroids directly in the microfluidics chip via automated widefield fluorescence microscopy. We were able to successfully measure drug-induced cytotoxicity in tumor spheroids, but had difficulties extending the system to measure T cell-mediated tumor killing.

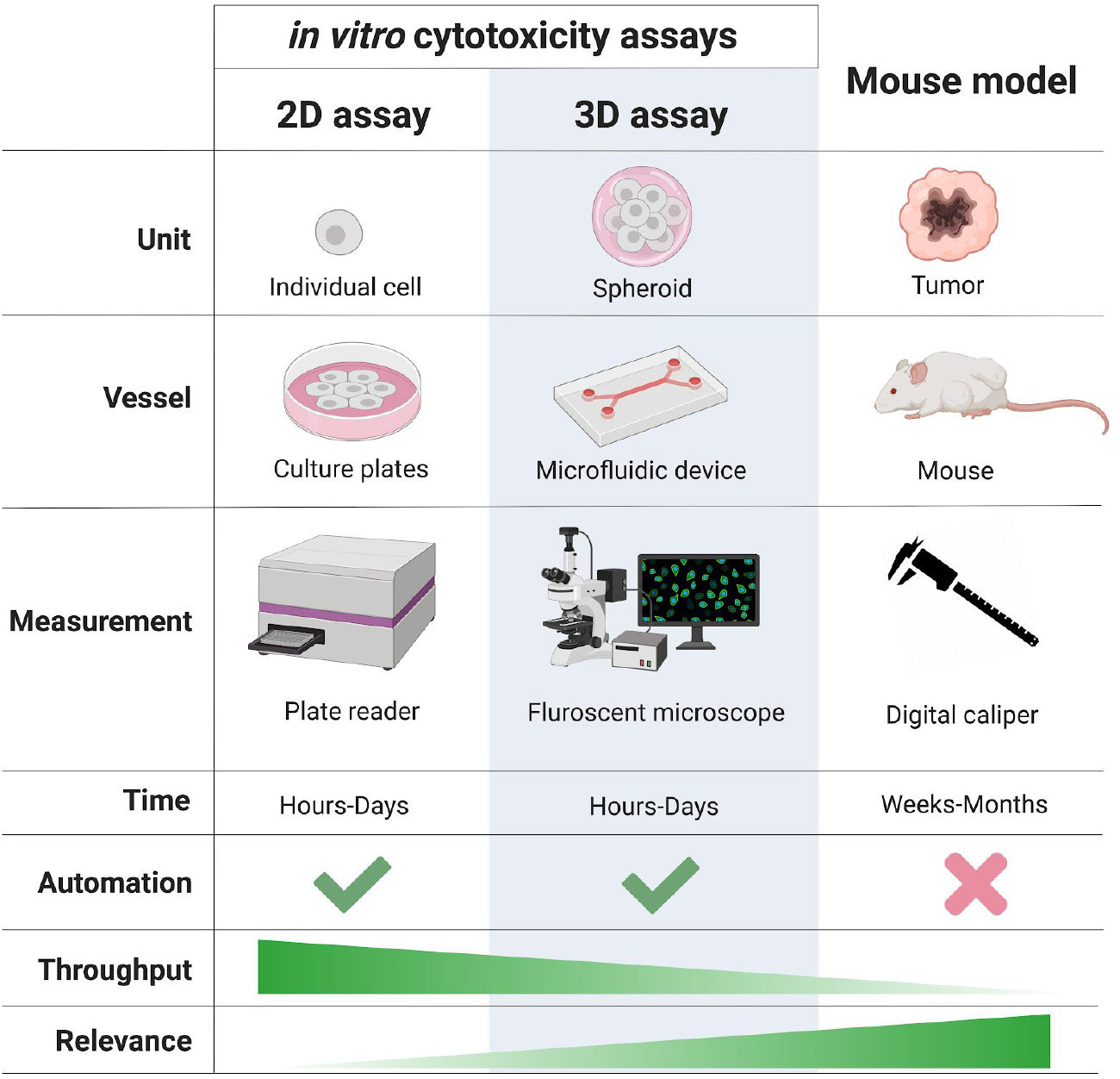

## Introduction

One area of interest in immunotherapy research is to modify cytotoxic T cells to make them better and more efficient at killing cancer cells (Ho et al. 2003; June 2007; Aksoy et al. 2019). Once T cells have been modified, their cytotoxic activity needs to be measured. T cell-mediated cytotoxicity assays have mainly been restricted to either two-dimensional (2D) *in vitro* studies or *in vivo* mouse models. 2D *in vitro* assays have the advantage of being high throughput and relatively cheap and fast compared to *in vivo* mouse studies, however the conditions are considered to be so artificial that the results do not always translate well to *in vivo.* Our goal was to explore the potential of an *in vitro* 3-dimensional (3D) cell culture system to study T cell-mediated cytotoxicity, and whether such a system could be a feasible intermediate between 2D *in vitro* and *in vivo* assays. Our thought was that a 3D T cell-mediated cytotoxicity assay has the potential to allow researchers to more rapidly and relevantly test the effect of T cell manipulation on T cell cytotoxicity before moving to the mouse model.

We had a few prerequisites for our system: 1) T cells must be added to multicellular tumor spheroids already suspended in a gel matrix so that the T cells would be required to migrate to the spheroids, 2) minimal manipulation of the samples post-coculture (which led us to explore an image-based assay for cytotoxicity), 3) larger sample size than confocal imaging provides (more spheroids being imaged per sample via widefield microscopy). We came up against a variety of challenges when exploring this system, which we discuss in this manuscript.

Why culture cells in 3D? Traditionally, *in vitro* experiments are conducted in 2D in a vessel, such as a dish or flask, and with liquid culture media. Under these conditions, adherent cell types will sink, adhere to the bottom of the vessel and grow as a monolayer, and suspension cells float in the liquid culture media. 2D culture conditions are artificial for adherent cell types because of limited cell to cell contact, contact with an artificial surface (the vessel) on the underside of the cells, and exposure to culture media on the upper surface of the cells. While suspension cells can form clusters suspended in liquid media in 2D culture conditions, there is no matrix for the cells to adhere to in order to move with directionality. These inherent characteristics of 2D cell culture can alter gene expression, in turn altering cell phenotype and behaviour (Edmondson et al. 2014; Knowles and Phillips 2001). Many cells can be forced to grow into aggregates, often called multicellular spheroids, by depriving them of a surface to grow on. This can be accomplished by various methods including using ultra-low attachment vessels, seeding a cell suspension in upside down droplets of media (hanging-drops), and u-bottom multiwell plates. 3D cell culture conditions usually include a gel matrix that the cells are suspended in and can interact with (although aggregates of cells grown in liquid media are often also referred to as 3D cell culture). Residing in a gel, cells can form cell-cell contacts across the entire cell surface, can retain or form multicellular aggregates, and can secrete extracellular matrix (ECM) and other factors to create and modulate a microenvironment. Furthermore, the gel provides a scaffold for cells to migrate through in 3D space. 3D cell culture conditions are thought to be somewhat closer to *in vivo* conditions than 2D culture (Edmondson et al. 2014).

In the context of immunotherapy research, *in vitro* studies of exogenously-derived T cell-mediated cytotoxicity of cancer cells have mainly been conducted in 2D culture. 2D conditions are particularly artificial in this system because the cancer cells are on the bottom of the vessel and the T cells have immediate access by function of gravity to come into contact with their targets instead of being required to migrate towards them. Additionally, the difference in cancer cell phenotype grown in a monolayer versus in a 3D can affect how well the T cells can target the cancer cells. One study found that tumor cells cultured as multicellular spheroids (in liquid media, not a gel matrix) did not activate cytotoxic T cells as efficiently as the same cells grown in a 2D monolayer due to defective antigen presentation (Dangles-Marie et al. 2003).

Recently, 3D cell culture systems have been explored in immunotherapy research, with studies using material from patient or mouse tumors to generate spheroids that retain tumor architecture (Jenkins et al. 2018; Deng et al. 2018; Pavesi et al. 2017; Courau et al. 2019). Pavesi et al. developed a 3D cell culture system with a microfluidics chip to study the cytotoxic effect of engineered T cells on cancer cells (Pavesi et al. 2017). In their system, single cancer cells or multicellular spheroids were seeded in a collagen gel in the center channels of the microdevice and T cells were added to a side channel. The T cells had to migrate into the gel to kill the cancer cells and cell death was monitored using confocal microscopy. They found that T cells only migrated into the gel when the cognate tumor cells were present, and that the T cells were more efficient at killing cancer cells in 2D culture versus in 3D culture. Jenkins et al. adopted the same 3D culture system to study the effect of anti-PD1 therapy with mouse or patient derived-organotypic spheroids (Jenkins et al. 2018). In this system, the immune cells in the spheroids are tumor infiltrating lymphocytes (TILs) that had already infiltrated the tumor *in vivo* in the mouse or patient. They used the microfluidics device to test the effect of anti-PD1 therapy on the antitumor activity of the TILs in the spheroids being cultured in a collagen gel. In another study, Courau et al. co-cultured T and NK cells with multicellular tumor spheroids (in liquid media) to study the characteristics of the immune cells that successfully infiltrated spheroids (Courau et al. 2019). Immune cells that had infiltrated spheroids were compared to immune cells that had not infiltrated via flow cytometry analysis. The volume of spheroids was measured using brightfield microscope images and caspase 3/7 activation via live fluorescence widefield microscopy to determine cell death in the spheroids. We wondered if we could adapt the 3D coculture system described by Pavesi et al. and Jenkins et al. to study exogenous T cells, that can be manipulated beforehand, and observe their cytotoxic effect on multicellular tumor spheroids being cultured in collagen gel, but to increase the sample size of spheroids measured via automated widefield fluorescence microscopy.

## Results

### B16-F10 cells can be killed by pmel-1 T cells *in vitro*, but inconsistently

We wanted to start with a model system that has T cells with a known target. We started with the well-characterized murine melanoma model pmel-1, which is used for immunotherapy studies on adoptive cell therapy (ACT) treatment of malignant melanomas (Overwijk et al. 2003). The CD8 T cells in the pmel-1 transgenic mouse express the T cell receptor (TCR) specific for the gp100 antigen that is presented by B16 melanoma cells. *In vivo* studies have demonstrated that ACT using pmel-1 T cells slows the growth of B16 melanoma tumors in mice (Overwijk et al. 2003; Abad et al. 2008).

To begin, we wanted to measure pmel-1 T cell-mediated killing of B16-F10 cells *in vitro* in 2D cell culture. After trying various approaches to measure T cell-mediated killing, we settled on a lactate dehydrogenase (LDH) release-based assay (CytoTox-ONE Homogeneous Membrane Integrity Assay, Promega) (Riss et al. 2019). Dying cells have damaged cell membranes causing cytosolic components, such as LDH, to leak into the culture media. The CytoTox-ONE assay uses LDH in the culture media to convert resazurin to fluorescent resofurin. The resulting fluorescent signal is proportional to the number of non-viable cells in a sample. An advantage to this plate-based assay is that the cancer cells do not need to be removed from the wells to measure cytotoxicity, which is especially convenient with adherent cell types that would require trypsinization. However, the assay does produce noisy data and a set of control wells with T cells alone is required to subtract background, overall increasing the number of replicates that are necessary.

To start, we used pmel-1 T cells that were day 4 and day 11 post-activation with the hgp100 peptide to co-culture with B16-F10 cells. We co-cultured the cells overnight at effector to target ratios of 4:1, 2:1, 1:1, and 1:2, and with or without pretreating the B16-F10 cells with interferon gamma (IFNγ). B16-F10 cells are known to have low presentation/expression of MHC Class I (MHC I) and IFNγ has been shown to upregulate MHC I on these cells (van den Boorn et al. 2010), which can further sensitize them to targeting by T cells. We saw only very minimal B16-F10 cell death when co-cultured overnight with day 4 T cells regardless of IFNγ treatment (see Supplemental Figure 1a). We saw an increase in cytotoxicity with the day 11 T cells co-cultured with B16-F10 cells (Supplemental Figure 1b). In this experiment, 80 to 90% of the B16-F10 cells were killed with ratios of T cells to target cells of 4:1, 2:1, and 1:1, and there was a decrease to 60% cytotoxicity at a 1:2 ratio. However, IFNγ pretreatment of the B16-F10 cells was not necessary for the day 11 T cells to target them. After washing the T cells out of the well plates, it was clear that the B16-F10 cells had been killed (example brightfield images are in Supplemental Figure 1c).

While we were exploring different methods to measure T cell-mediated cytotoxicity in 2D cell culture, we noticed phenotypic changes in the B16-F10 cells that were being maintained in culture. Most strikingly, the B16-F10 cells visibly appeared to stop producing melanin after being in culture for several weeks (see Supplemental Figure 2 for light microscope images to compare early and late passage number cells). The longer the cells had been in culture, the less melanin they appeared to produce, and the culture media would remain the pink from the phenol red (instead of turning brown as it does with cells that are producing melanin). Pmel is a structural protein necessary for the formation of melanosomes (Theos et al. 2005). We thought that if there was less melanin production in the cells, perhaps there was less expression of the Pmel protein, and therefore less antigen for the B16-F10 cells to be targeted by the pmel-1 T cells. We repeated the co-culture experiments using both early passage B16-F10 cells (that were still producing melanin) and late passage number B16-F10 cells. We used pmel-1 T cells that were 4, 7, and 11 days post-activation with the hgp100 peptide. The results from this experiment suggested that the passage number of the B16-F10 cells affected the extent to which they were targeted by the pmel-1 T cells (Supplemental Figure 3). At a 4:1 ratio of effectors to targets, the day 4 T cells were able to kill almost 100% of the early passage B16-F10 cells, but only when the cancer cells had been pretreated with IFNγ (Supplemental Figure 3a). The day 7 T cells could kill either passage number of B16-F10 cells, but only with IFNγ pretreatment of the cancer cells (Supplemental Figure 3b). And, as we saw in our first experiment, the day 11 T cells were able to kill either passage number regardless of IFNγ pretreatment (Supplemental Figure 3e and f). Flow cytometry profiles of the T cells at different days post-activation are in Supplemental Figure 4.

The results suggested that IFNγ pretreatment of the melanin-producing (early passage number) B16-F10 cells, but not the late passage B16-F10 cells, sensitized them to targeting by the day 4 pmel T cells. We stained both passage number B16-F10 cells, with or without IFNγ treatment, with antibodies to detect surface MHC I expression via flow cytometry (Supplemental Figure 5). IFNγ treatment induced surface MHC I expression in both the early and late passage number B16-F10 cells, however MHC I levels were higher in the early passage B16-F10 cells compared to the late passage cells. We also checked PD-L1 levels on the cells and found a similar pattern of expression as MHC I between the passage numbers.

To repeat the above experiments we ordered another batch of B16-F10 cells from the ATCC (Manassass, VA) to have more “early passage” cells to work with. The cells do not arrive with a passage number, we were simply assigning the cells “passage 0” when we started culturing them. When we first thawed the new batch of cells they were not producing melanin, but after 2-3 passages melanin deposits appeared. By passage 5 they appeared similar to the early passage cells from the first batch we ordered (Supplemental Figure 6).

However, when we repeated the experiments two more times, there was no clear indication that the differing phenotypes of the B16-F10 cells affected how they were targeted by the pmel-1 cells that were 4, 7, or 11 days post-activation with the hgp100 peptide (see all results in Supplemental Figures 7 and 8).

### B16-F10 cells can be replaced with hgp100-pulsed MC38 cells

Because of the variability with our results co-culturing the pmel-1 T cells and B16-F10 cells, we tried using a murine colon cancer cell line, MC38, and sensitized them to pmel-1 T cells by pulsing with the hgp100 peptide. Pulsing the MC38 cells with the hgp100 peptide (the cognate antigen for the pmel TCR) allowed for the MC38 cells to be killed by the pmel T cells (Figure 1). First, we used day 11 pmel T cells to co-culture with MC38s that had been pulsed for 1 hour with varying concentrations of peptide (Figure 1A). We saw that there was close to 100% killing at a 4:1 ratio of T cells to cancer cells, and that the cytotoxicity decreased as the ratio of T cells to cancer cells decreased. The different concentrations of peptide did not affect how much the MC38 cells were killed. There were only background levels of cytotoxicity without the peptide pulsation, even at the highest ratio of T cells to target cells. When we repeated the experiment we wanted to see if day 4 (post-activation) pmel T cells would kill the pulsed MC38 cells and if the MC38s could still be targeted if the T cells were added the day after the addition of peptide (Figure 1B). Day 4 T cells were just as effective as the day 11 T cells at killing the pulsed MC38s if they were added to the culture 1 hour after peptide pulsation. However, the effect of peptide pulsing had worn off almost completely when the T cells were added 24 hours after the peptide had been added. Addition of peptide alone was not toxic to the MC38 cells (Figure 1c). Not only were the hgp100-pulsed MC38s killed at comparable levels to B16-F10 cells, if not more efficiently, but they were also more reliable to kill with pmel T cells in the 2D cytotoxicity assay without the complications of passage number and IFNγ pretreatment.

**Figure 1:**
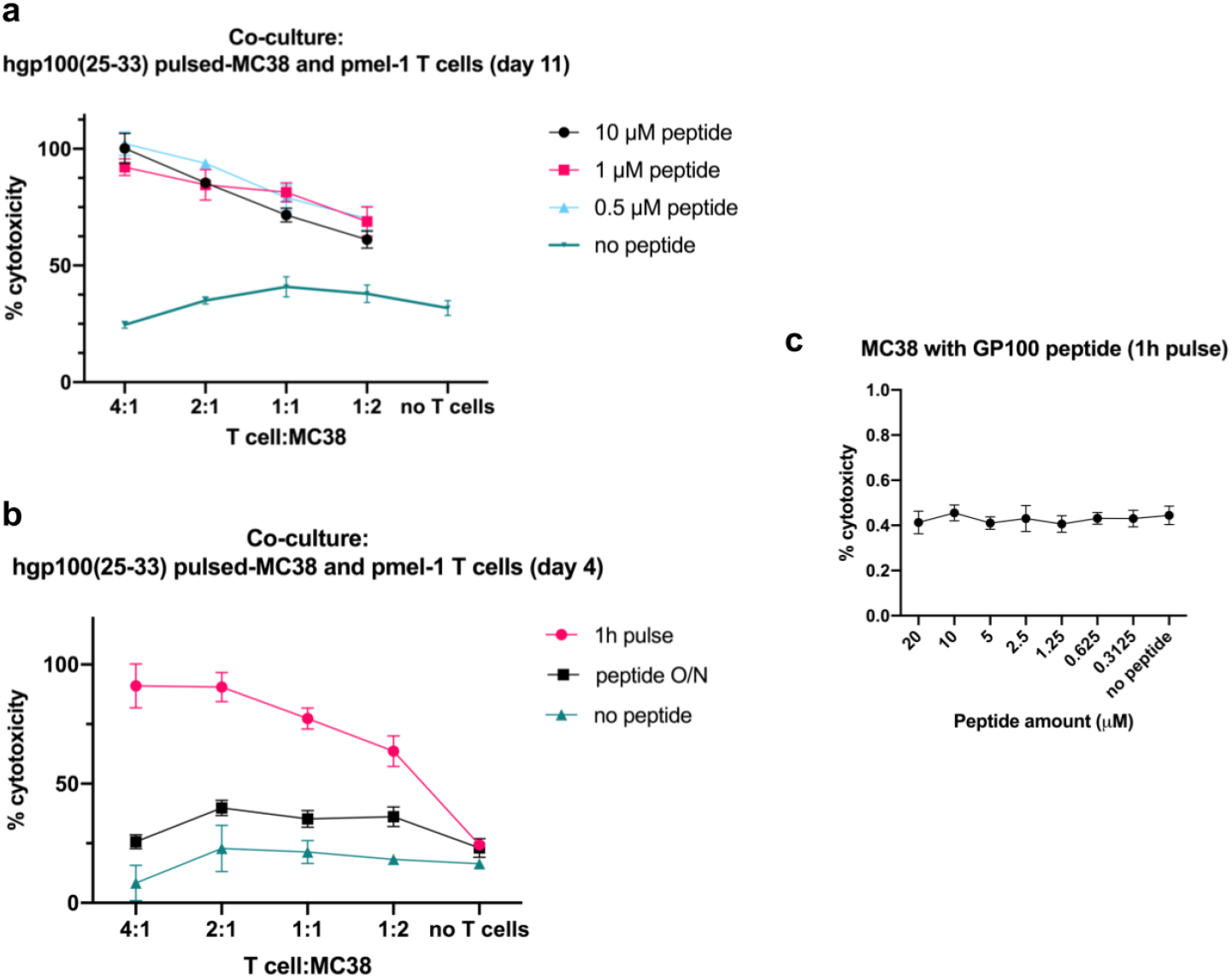
Pmel-1 T cells can kill MC38 cells that have been pulsed with the hgp100 peptide. **a)** MC38 cells adhered to the bottom of 96 well plates were pulsed for one hour with hgp100 peptide at the noted concentrations before co-culture with Pmel-1 T cells (at day 11 post-activation with hgp100 peptide). **b)** MC38 cells were pulsed for one hour or overnight (O/N) with 1 µM hgp100 peptide before co-culture with pmel-1 T cells (at day 4 post-activation with hgp100 peptide). 24 hours after the start of co-culture. **c)** MC38 cells were pulsed with the hgp100 peptide at the indicated concentrations for 1 hour and then allowed to grow overnight. **a-c)** LDH release was measured with a plate reader using the CytoToxONE assay (Promega).

### MC38 cells form multicellular spheroids, can be grown, treated, and assayed for cell death in 3D culture within microfluidics chips

Our aim was to transition to exploring a 3D setup for T cell-mediated cytotoxic studies once we had verified T cell-mediated killing in the 2D assay. Therefore, while we were working on the 2D cytotoxicity assay with the pmel-1/B16-F10 model, we were also attempting to grow tumor spheroids with various murine cancer cell lines. We used the simple, scaffold-free, “hanging drop” method to experiment with generating tumor spheroids from cancer cell lines (details in the Supplemental Materials and Methods) (Müller and Kulms 2018). Briefly, a single cell suspension was pipetted in 20 ul droplets on the lid to a 10 cm culture dish and inverted. The droplets were incubated and monitored daily under the microscope for cell aggregation.

B16-F10 cells did not form spheroids easily, requiring about 1-2 weeks before the cell aggregates could withstand being passed through nylon mesh filters with pore sizes of 40 and 100 um. Luckily, MC38 cells formed spheroids easily and reliably. After 24 hours, a single cell suspension of MC38 cells consistently formed spheroids that were spherically shaped, uniformly sized, and withstood size filtering (Figure 2a). Furthermore, for eventual image-based quantification of cell death in 3D culture using live/dead fluorescent staining, MC38 spheroids were more straightforward than B16-F10 spheroids. As the B16-F10 cells formed spheroids they appeared to produce melanin, an observation recently reported (Chung, Lim, and Lee 2019). When we stained B16-F10 spheroids in collagen gel with AO/PI live/dead stain, the melanin deposits in the cells masked the fluorescent signal from the stain, making quantification of the fluorescent signals difficult (Supplemental Figure 9).

**Figure 2:**
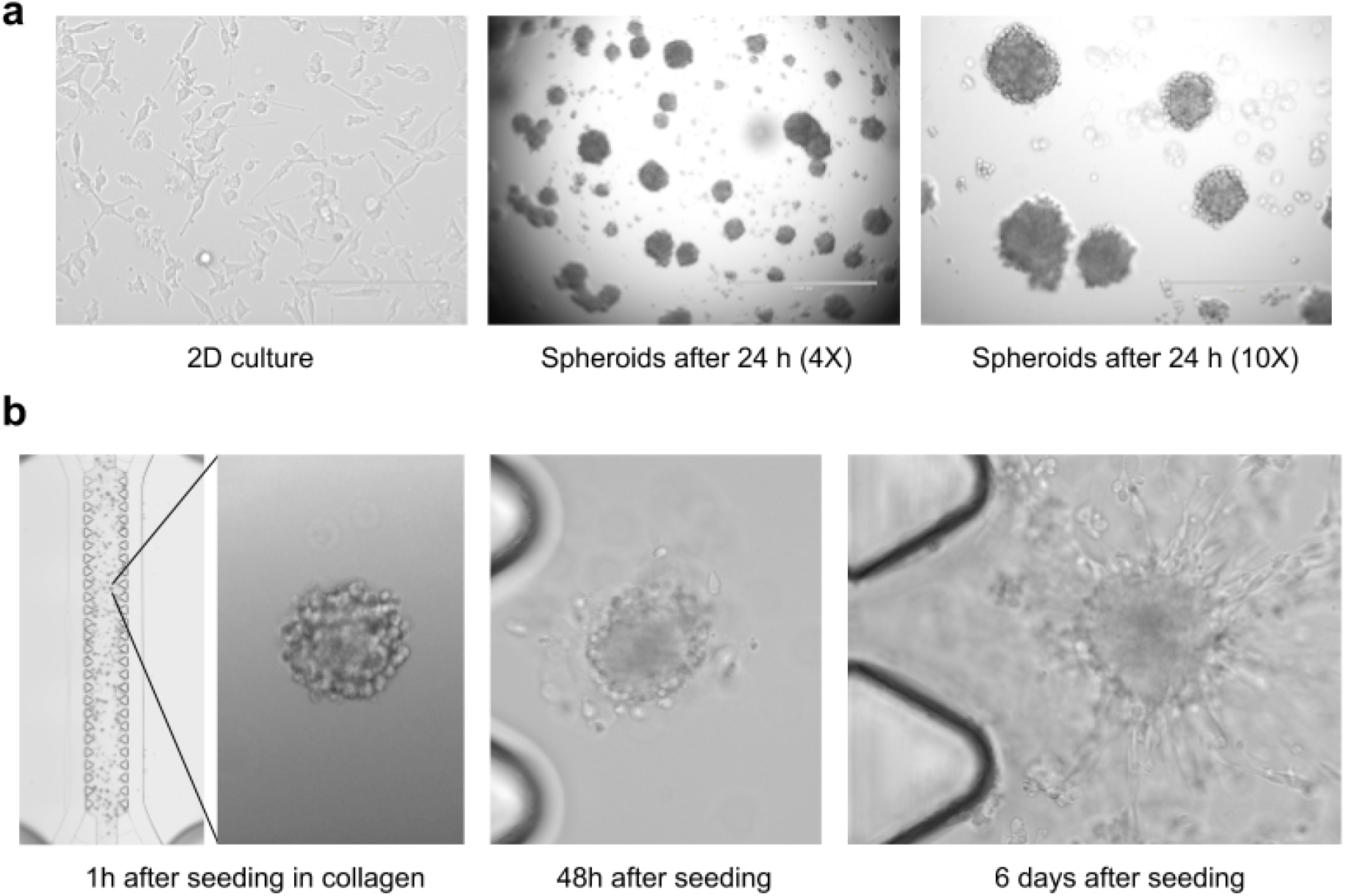
**a) MC38 cells form spheroids after 24 hours in culture.** MC38 cells in a monolayer in 2D cell culture (left), MC38 spheroids after 24 hours of culture in hanging droplets shown in a single hanging droplet at 4X (middle) and at 10X (right). Scale bars are at 200 µm, 1000 µm, and 400 µm for the left, middle, and right images respectively. Cultures were imaged on an EVOS microscope. **b) MC38 spheroids grow in collagen within a 3D cell culture chip (AIM Biotech).** A view of a whole channel in the 3D culture chip with MC38 spheroids seeded in a collagen hydrogel for 1 hour after being formed in hanging droplets overnight (left). A single spheroid is enlarged to the right. An MC38 spheroid that was cultured for 48 hours in the 3D cell culture chip (middle) and after 6 days (right). Samples were imaged on the Keyence BZ-X710 with a 2X objective (left) and 20X objective (insert, middle, and right).

Based on the protocol described by Jenkins et al., we harvested the MC38 spheroids from the hanging droplets, size filtered them by sequentially passing them through 40 um and 100 um nylon mesh filters, then resuspended them in a collagen gel mixture, and pipetted the spheroid/gel mix into the central channel of microfluidics chips (Jenkins et al. 2018). The gel mixture was allowed to polymerize for 30 minutes at 37 °C before the culture media, treatments, or T cells were added to the side channels. Because for our eventual co-culture experiments with T cells we weren’t sure how long we would need to culture the T cells with the spheroids for and how well the spheroids would stand up to extended 3D culture we tried culturing the MC38 spheroids in the collagen gel in the microfluidics chips for 6 days. The spheroids thrived and starting growing projections of cells out from the main body (Figure 2b).

To eventually measure the extent of T cell-mediated killing of tumor spheroids in co-culture, we required a reliable method to quantify cell death in the spheroids. We decided to take an image-based approach because we wanted to manipulate the samples as little as possible after co-culture in the chips. The extraction of spheroids from the collagen gel for downstream analysis could result in increased cell death and loss of sample, and would add extra steps and time to the co-culture protocol. Our goal was to image many spheroids at once across each of the entire central channels in the microchip to increase sample size and reduce sampling bias. We chose not to use confocal microscopy to image the spheroids because of the low number of spheroids that would be possible to image per sample and the potential for bias when selecting a spheroid to image. When we started our imaging experiments, we observed variable staining of spheroids dependent on where the spheroid was located in the central gel channel (with both Hoechst and the dead cell stain we were using). If only spheroids in the center of the gel channel were imaged, the dead cell signal would be lower than if spheroids towards the side channels were imaged. With our setup we could image all the spheroids in a channel (about 50-125 spheroids per channel), in 3 channels per chip. We accomplished this by imaging with the Keyence BZ-X710 system, which is a widefield fluorescence microscope with automation. To image one central gel/spheroid containing channel with a 20X objective and 20 μm step-size required a 3×11 grid and a stack of 6-9 images per field of view. To measure cell death in spheroids, we used SYTOX red dead cell stain (Thermofisher), in combination with Hoechst (to stain all nuclei). The SYTOX dye can only permeate the cell surface of dead cells and, once inside, the dye binds to DNA and fluoresces.

To develop a quantification method using the dead signal in our images, we generated a test dataset by treating MC38 spheroids cultured in collagen in 3D cell culture chips with a titration of staurosporine (an apoptosis-inducing antibiotic (Belmokhtar, Hillion, and Ségal-Bendirdjian 2001)). We compared the viability of MC38s in 2D culture treated with a titration of staurosporine to the percent live in 3D cell culture culture (Figure 3a and b). Representative images for three concentrations of staurosporine are shown in Figure 3b. The whole channel of spheroids is shown on the left (a stitched image). The blue square indicates single spheroids that have been expanded and are shown on the right.

**Figure 3:**
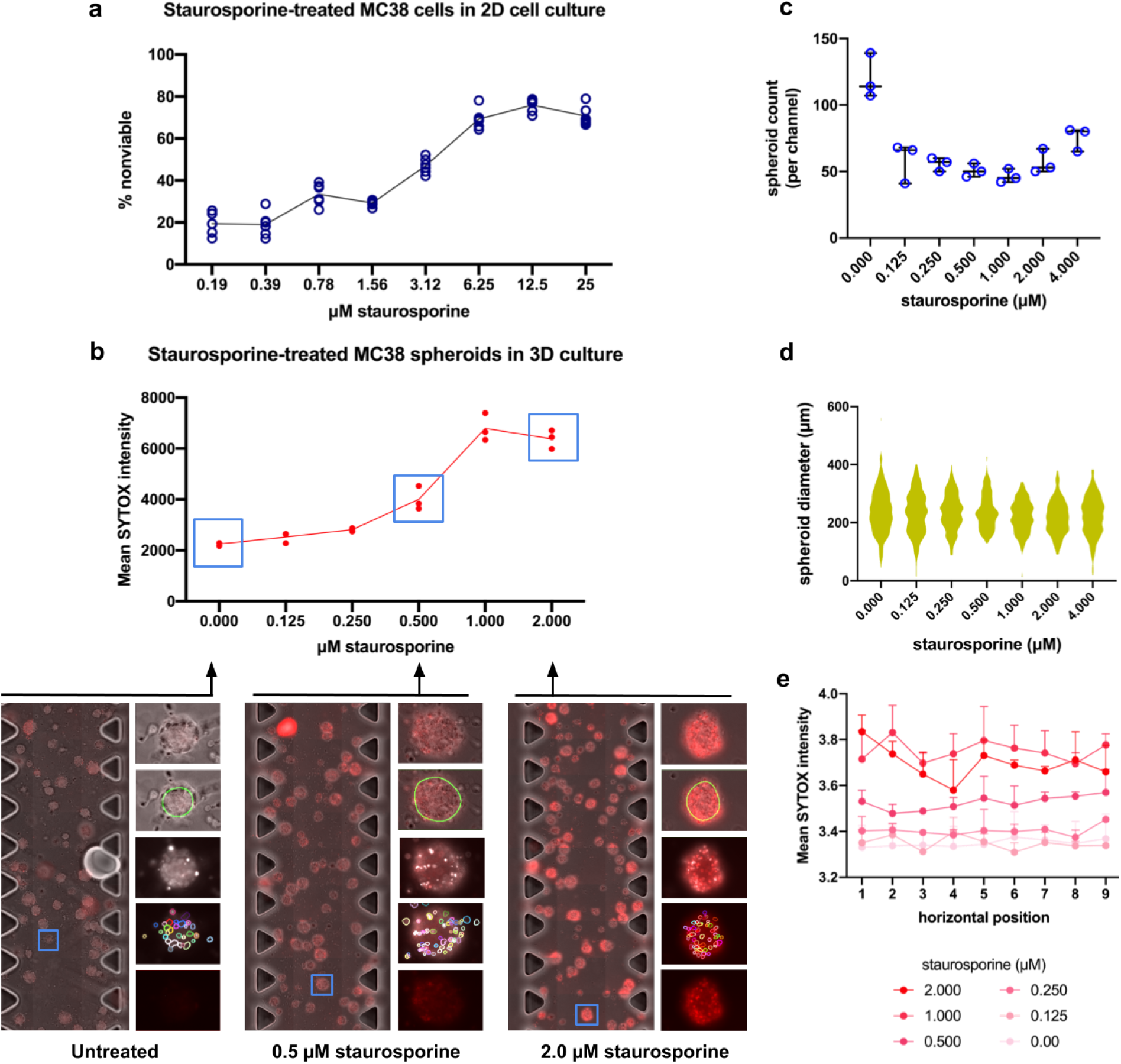
The concentration-dependent cytotoxic effect of staurosporine on MC38 spheroids in 3D culture can be measured from microscope images. **a)** MC38 cells grown in 96 well flat bottom plates (2D cell culture) were treated with increasing concentrations of staurosporine for 24 hours. Alamar Blue was added and the cells were cultured for another 24 hours. The absorbance of the wells was measured on a plate reader and indicates the percent of viable cells in each condition. **b)** Collagen gel containing MC38 spheroids was seeded into 3D cell culture chips (AIM Biotech) and treated with the indicated concentrations of staurosporine in complete culture media for 24 hours. The samples were stained with Hoechst and SYTOX red dead cell stain in the culture chips for 30 minutes before imaging. The entirety of each channel within each chip was imaged with a 20x objective, 20 μm z-step, and a 3×11 grid using automated microscopy on the Keyence BZ-X710. Three replicates (three channels) were imaged for each condition and analyzed for SYTOX signal intensity. The quantification shown in the graph demonstrates the increase in SYTOX intensity as the concentration of staurosporine increases. Representative images are shown below the graph for untreated, 0.5 μM, and 2.0 μM staurosporine-treated samples with fixed SYTOX channel contrast (0-16000). A stitched image of the whole channel is shown at the left. The blue squares indicate the area that has been expanded to show a single spheroid on the right of each channel. From top to bottom: the layered image of the spheroid (with brightfield, Hoechst, and SYTOX), the spheroid boundary (green line) demonstrating object recognition of a spheroid, the layered image with only Hoechst and SYTOX channels, the object recognition of single cells in a spheroid (shown with multicolored lines), and the SYTOX channel alone. **c)** Spheroid counts per condition. The dots indicate the three replicates in each condition. **d)** Mean spheroid size and standard deviation. Between 140 and 360 spheroids were measured for each condition. **e)** The mean SYTOX intensity at horizontal positions across the spheroid-containing channel. Error bars indicate standard deviation.

Our primary approach for quantifying cytotoxicity within images of the 3D cultures required first identifying spheroid objects (illustrated by the green line drawn around spheroids in Figure 3b). These objects were identified within 2D brightfield images, first max-projected across the z dimension, using Ilastik (Berg et al. 2019) pixel classification models. Spheroid masks were derived from a voronoi-based segmentation (Jones, Carpenter, and Golland 2005) of the brightfield images using seed points calculated as local maxima in the distance transform image of a hysteresis threshold applied to pixel probabilities supplied by the model. Quantification of cytotoxicity for an individual spheroid was defined as the integration of SYTOX channel intensity over the 3D volume of the spheroid constrained only by the 2D boundaries determined from the max-z projection image. More details on this approach, as well as comparable approaches employed, are provided in Materials and Methods and Supplemental Figure 10. Our image analysis also included a spheroid count per channel and measurements of spheroid diameters (Figure 3c and 3d, respectively). The spheroid count was consistently higher in the untreated condition compared to any treatment (and this trend continued when we moved to co-culturing with T cells, shown in Supplemental Figure 12).

Through generating this test dataset, we learned that the time the SYTOX stain is in the chips affects how well the stain diffuses across the channel, which influences the SYTOX intensity. Therefore, we found that it was necessary to keep the staining time the same for each chip, which required staining the samples one at a time to account for the time it took to image a chip on the microscope. We analyzed the mean SYTOX intensities at horizontal positions across the channel for each of the concentrations and found them to be consistent horizontally across the channel for this experiment (Shown in Figure 3e). The results from this experiment gave us confidence that we could detect and measure changes in the extent of cell death from images of spheroids cultured in 3D.

### T cells require the presence of cytokines to induce migration into collagen gel

Our plan for co-culturing the MC38 spheroids with T cells in 3D culture was to have the spheroids in the gel in the center channel of the chip and to add the T cells to the side channels. Our assumption was that the activated T cells would migrate into the gel from the side channel and come in contact with the tumor spheroids. However, we found that T cells added to the side channel of a chip did not readily migrate into the collagen gel in the center of the chip. A recent preprint reported that the addition of the chemokine CXCL12 (ligand for the receptor CXCR4) allowed human T cells to migrate in a collagen gel (Abu-Shah et al. 2019). CXCR4 is constitutively expressed on T cells (Contento et al. 2008), and another receptor, CXCR3, is only expressed upon activation. Both receptors allow T cells to traffic to sites of inflammation, where their inflammatory cytokine ligands are being secreted (Hu et al. 2011; Qin et al. 1998). Because ligands for CXCR3 and CXCR4 are known to attract T cells and induce their migration, we tried adding them to our system to draw the T cells into the gel. We tested CXCL9, 10, 11 (ligands for CXCR3) and CXCL12 and started with a concentration of 0.3 ug/ml based on the preprint mentioned above (Abu-Shah et al. 2019). We tried adding the cytokine either to the opposite side channel from the T cells or directly into the collagen gel mixture before loading it into the central channel. We did the experiment once and observed the most migration with CXCL11 added directly to the collagen gel. We continued with a titration of CXCL11 in the gel and found there was the most migration with the highest concentration of CXCL11 used, which was 2.7 ug/ml (shown in Figure 4, brightfield on the left and fluorescently stained T cells on the right). Because of limited access to mouse splenocytes, we used human healthy donor T cells (pan T cells isolated from PBMCs) and human cytokines for these experiments. We verified that murine CXCL11 worked to attract pmel-1 T cells into the collagen gel, although we saw less migration compared to the human T cells (overall the human T cells appeared to thrive more in the 3D system).

**Figure 4:**
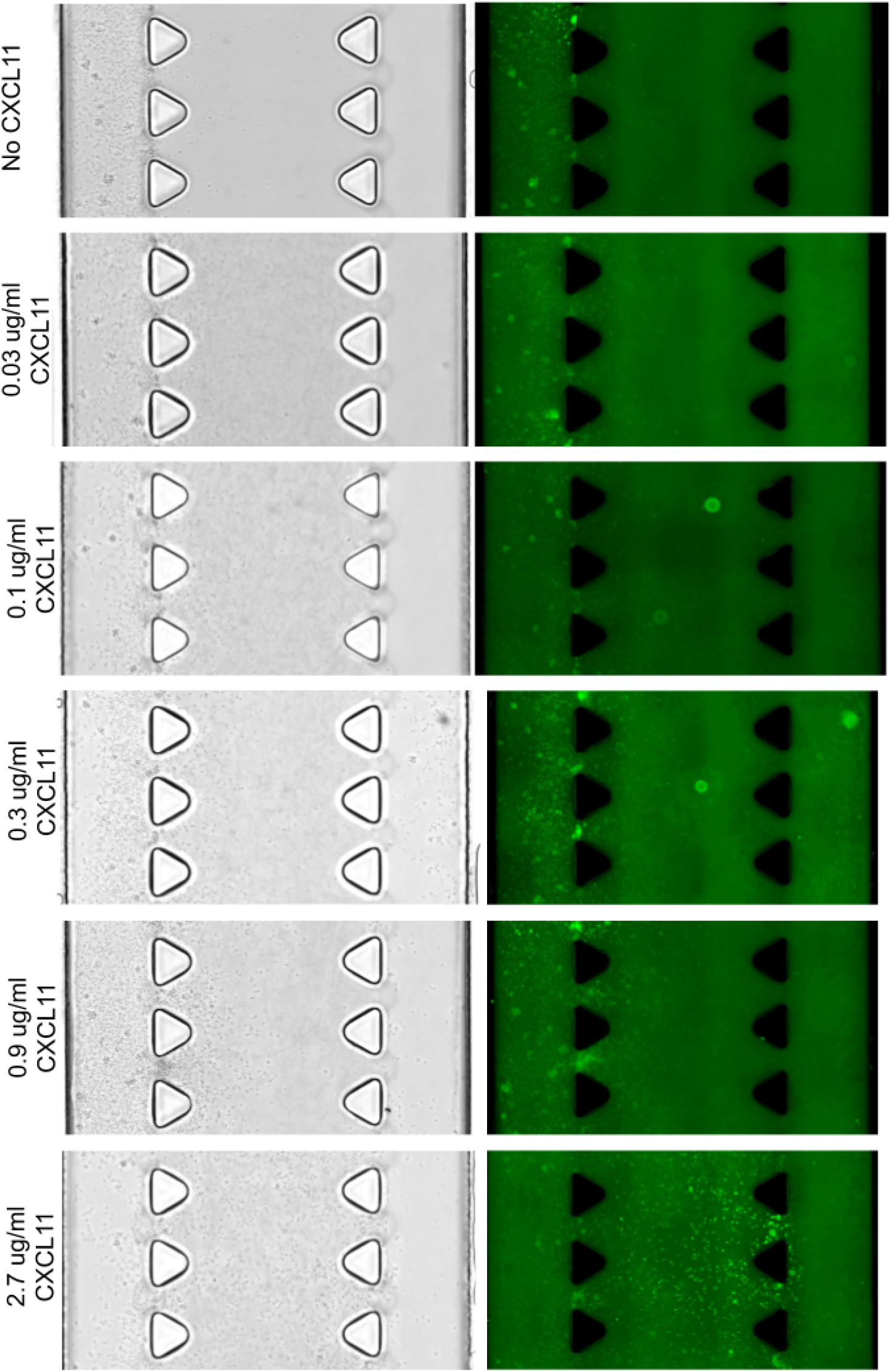
T cells migrate into collagen gel containing the CXCR3 ligand CXCL11. Collagen gel with titrated amounts of CXCL11 was pipetted into 3D cell culture chips (AIM Biotech) and allowed to polymerize. Activated healthy human donor T cells were stained with IncuCyte CytoLight Rapid Green Reagent (Essen Bioscience) and added to the left side channel in the chips and incubated overnight. The entire of the three central channels in each chip was imaged on the Keyence BZ-X710 with a 20X objective, 20 μm z-step, and automated microscopy.

### MC38 spheroids can present peptide in 3D cell culture

Another component to co-culturing the MC38 tumor spheroids with pmel T cells was the transitory nature of the hgp100 peptide presentation. As shown in Figure 1, the MC38s were only effectively targeted by the pmel T cells when the T cells were added 1 hour after pulsing the MC38s with the peptide, but not when the T cells were added one day after peptide addition. As the spheroid formation for MC38 cells requires one day, it was not feasible to pulse the MC38 cells before allowing them to form spheroids. At first, we thought to transiently express the pmel protein in MC38 cells via electroporation. We optimized the electroporation settings for MC38 cells using a GFP-encoding plasmid and found via western blotting (probing with an anti-gp100 antibody) that gp100 peaked at 24 hours post-electroporation and diminished significantly by 48 hours (Supplemental Figure 11). However, this approach became unnecessary when we became aware of an antibody specific for another peptide (SIINFEKL) only when it is bound to MHC1. If we added gp100 peptide to the already formed spheroids, we had no way to test whether the MC38 spheroids in the gel could take up the hgp100 peptide and present it. Therefore, we used the SIINFEKL to test the peptide addition and presentation in 3D culture. Using fluorescent microscopy, it was clear that the addition of peptide to 2D and 3D cultures resulted in presentation (red signal in Figure 5a and b). We tried both adding the peptide directly to the gel with the spheroids or adding the peptide to the culture media at the side channel. We found that spheroids presented the peptide in both methods after 2 hours of incubation with peptide. However, 24 hours later only the spheroids with peptide in the culture media were still presenting (Figure 5b).

**Figure 5:**
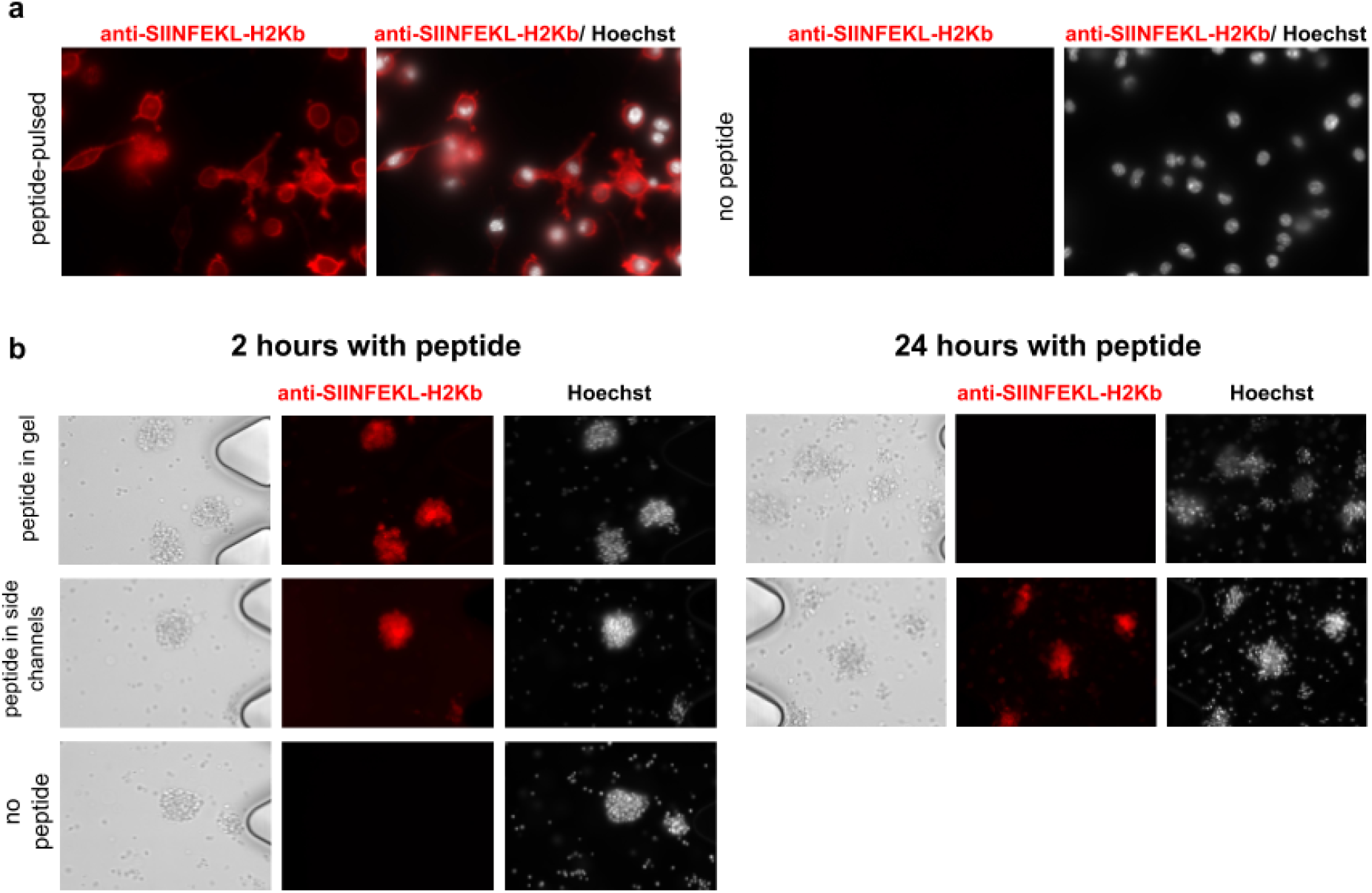
MC38 spheroids can present peptide when added to the gel or culture media. **a)** MC38 cells grown in 35mm culture dishes were pulsed with the SIINFEKL peptide (left panels) or unpulsed (right panels) before staining with an antibody against SIINFEKL bound to H2Kb (shown in red). Cell nuclei were stained with Hoechst. **b)** MC38 spheroids in collagen gel were seeded in 3D cell culture chips. The SIINFEKL peptide was added to the gel with the spheroids before seeding the chips (top row), only to the media in the side channels (middle row), or no peptide was added (bottom row). Sample were stained with the anti-SIINFEKL-bound H2Kb antibody (red) and Hoechst (white) and imaged at 2 hours post-peptide addition (left panels) and at 24 hours post-peptide addition (right panels). Imaging was performed on the Keyence BZ-X710 with a 20X objective.

### Pmel-1 cells can target MC38 spheroids within the gel in the presence of hgp100 peptide, however the results are unpredictable

At this point, we thought we had enough components of our system working to try co-culturing T cells with spheroids in 3D cell culture and measure T cell-mediated cytotoxicity (Figure 6). We used the same 3D cell culture chips (AIM Biotech). We seeded MC38 spheroids in collagen gel (containing CXCL11 and IL-2) into the central channels of the chips. Pmel-1 T cells at different concentrations were added to the left side channels once the gel/spheroid mix had polymerized in the central channels. Culture media without the hgp100 peptide or with the peptide (at 2 μM to try and compensate for the media being diluted across the collagen gel) was added into the right side channels. Co-culture proceeded for 24 hours before imaging. To verify that the T cells were cytotoxic, in parallel to each 3D co-culture experiment we conducted co-culture in 2D with the indicated ratios of T cells to MC38 cells, with hgp100 peptide (magenta) and no peptide (green) (Figure 6a, experiments 1-4). In all four 2D culture experiments we saw consistent killing of peptide-pulsed MC38 cells by pmel-1 T cells and the percent cytotoxicity was dependent on the ratio of T cells to MC38 cells.

**Figure 6:**
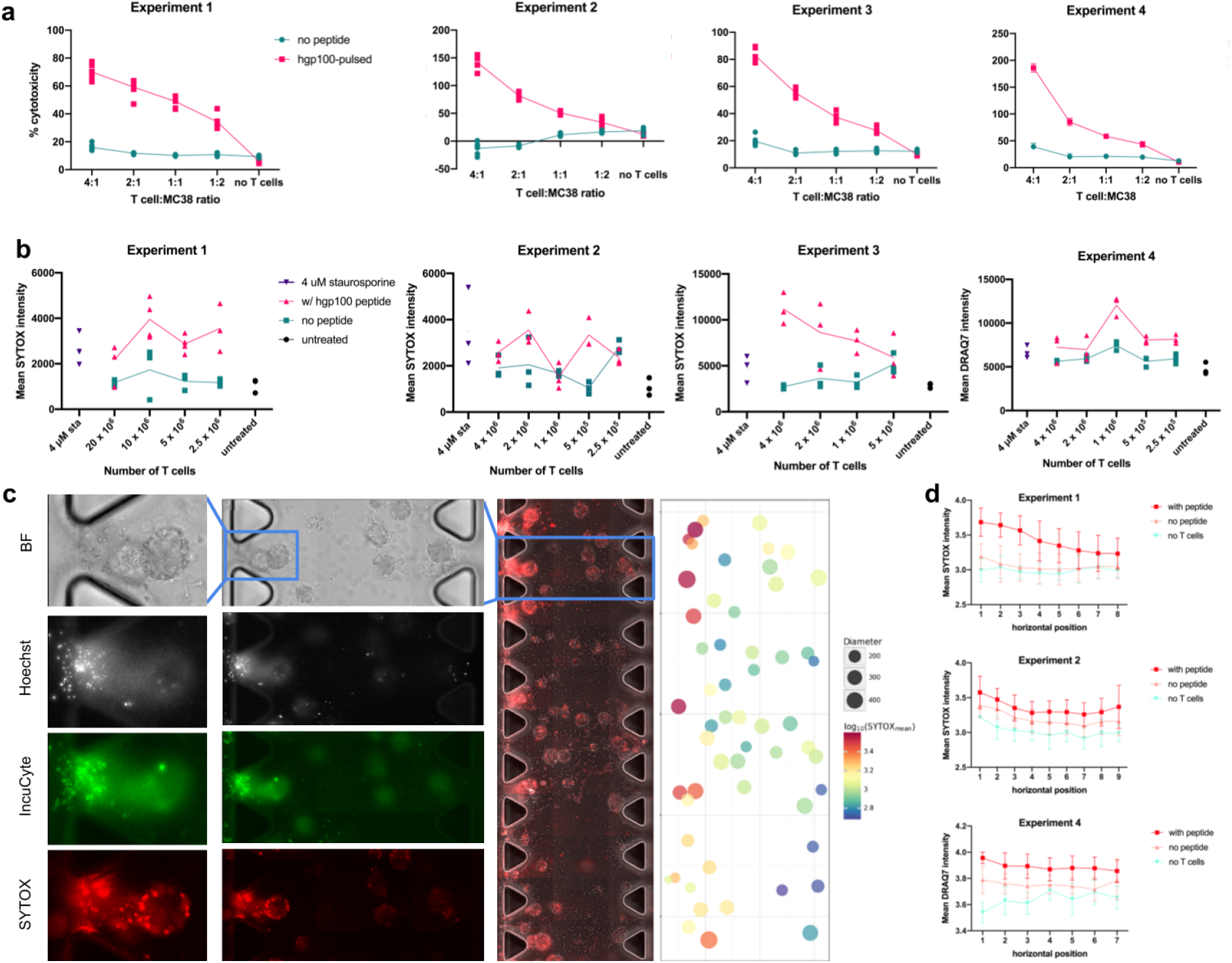
Measuring pmel-1 T cell-mediated cytotoxicity in 2D cell culture and 3D cell culture. **a) 2D co-culture:** MC38 cells growing in a monolayer in 96 well plates were either pulsed for 1 hour with 1 μM hgp100 peptide (magenta) or no peptide was added (green). Pmel-1 T cells were added at the indicated ratios to the MC38 cells and co-cultured for 24 hours. LDH release was measured with a plate reader using the CytoToxONE assay (Promega). Percent cytotoxicity was calculated as described in materials and methods. The results from four independent experiments are shown (experiment 1 - 4). **b) 3D co-culture:** MC38 spheroids in collagen gel were cultured in 3D cell culture chips with the hgp100 peptide (magenta) or without peptide (green) and the indicated numbers of pmel-1 T cells for 24 hours before staining and imaging. 4μM staurosporine (sta) was used as a positive control (purple) and untreated MC38 spheroids as a negative control (black). The entirety of each channel within each chip was imaged with a 20x objective, 20 μm z-step, and a 3×11 grid using automated microscopy on the Keyence BZ-X710. Three replicates (three channels on one chip) were imaged for each condition and analyzed for SYTOX signal intensity. Data points indicate replicates within each experiment. Experiments 1-3 used SYTOX as a dead cell stain. Experiment 4 used DRAQ7 as a dead cell stain. Mean SYTOX or DRAQ7 intensity was calculated with the “Spheroids” method, explained in the Materials and Methods and illustrated in **Supplemental Figure 10**) **c)** An example co-culture image (with hgp100 peptide) shows how T cells entering the channel on the left result in a spatial SYTOX intensity bias. A stitched image of the whole channel is shown in the center panel. The blue squares indicate the areas that have been expanded to the left, where the imaging channels have been separated out: from top to bottom: brightfield (BF), Hoechst, IncuCyte green (T cells only), and SYTOX red (dead cells). The scatterplot on the right shows the spheroids identified in the image in the middle panel scaled by inferred size and colored by mean SYTOX intensity (showing the decrease of signal from left to right). **d)** Mean SYTOX or DRAQ7 intensity for each condition horizontally across the central spheroid-containing gel channel. The mean intensity for each horizontal position in each replicate in each T cell count within each condition (no peptide or with peptide) was averaged and shown with standard deviation. From top to bottom: Experiment 1, Experiment 2, and Experiment 4. Conditions with the hgp100 peptide (red), no peptide (pink), no T cells (green).

The image analysis results from the 3D co-culture experiments are shown in Figure 6b. Experiment 1 was analyzed before the “Primary” method was optimized for object recognition of spheroids. We repeated the experiments three times using the SYTOX dye (Figure 6b Experiments 1-3) and one time with a different dead cell dye, DRAQ7 (Figure 6b). In each experiment, we treated one chip with 4 μM staurosporine as a positive control (we expected to see a high level of SYTOX (dead) signal in these samples) (Figure 6b, purple). As shown in the graphs in Figure 6b, the dead cell signal intensity for the co-culture conditions that included hgp100 peptide (magenta) were usually higher than the conditions that had no peptide in the culture (green). However, only one experiment (Experiment 3) showed a trend of higher SYTOX signal with increasing T cell numbers in the peptide-treated samples. The mean SYTOX intensity for the staurosporine-treated positive control was similar or even lower than the co-culture conditions that included peptide.

Example images from a co-culture experiment with peptide are shown in Figure 6c. The whole channel is shown as a stitched image in the center panel. The blue rectangles denote areas that were expanded to the panels shown with the imaging channels separated as follows: (from top to bottom) brightfield (BF), Hoechst (nuclei in all cells), IncuCyte dye (T cells only), and SYTOX (any dead cells). Note the T cells (green) moving into the gel from the left side channel. We included the IncuCyte dye in experiments 1 and 2 only to visualize the T cells, but eliminated the dye in experiments 3 and 4. Spheroid size did not change between conditions. Spheroid counts were higher in untreated samples than any of the treated, similar to the results with the staurosporine titration in Figure 3. Spheroid size and counts for the 3D co-culture experiments are shown in Supplemental Figure 12.

In our images, we noticed that the SYTOX signal appeared to be more intense on the left side of the central channel with the gel (the side where the T cells were entering the gel), as is shown in Figure 6c. We thought this was logical if the T cells are coming into contact with these spheroids first. We looked that the mean SYTOX intensity at horizontal positions across the central gel channel and saw that the signal was higher on the left side of the channel than the right in experiment 1 and less so in experiment 2 (Figure 6d). Unfortunately, the orientation of the chips in experiment 3 changed from condition to condition so we were unable to analyze the horizontal bias from that experiment. The SYTOX stain is added to the side channels and incubated for 30 minutes before imaging. We found that sometimes the short staining time would result in irregular staining of the channel (the spheroids at the edges of the channel tended to be stained more brightly than spheroids in the center of the channel). Increasing the staining time to 1 hour helped, but inconsistent staining still occurred sometimes and became cumbersome as the number of samples increased. Pavesi et al. used another dead cell stain, DRAQ7, to label dead cells in their 3D co-culture studies (Pavesi et al. 2017). DRAQ7 is a stable molecule and nontoxic to live cells, and can therefore be included in the culture media at the beginning of co-culture. We thought perhaps having the dead cell dye already mixed into the collagen with the spheroids before polymerization would result in more uniform staining. In Experiment 4 we added DRAQ7 to the collagen gel mixture before adding the spheroids and did not stain with SYTOX on the day of imaging. There appeared to be less of a horizontal bias to the side where the T cells enter the gel in the peptide-treated conditions (Figure 6d), but it is hard to say whether that was due to inefficient T cell-mediated killing in the 3D co-culture (Figure 6b).

## Discussion

In this manuscript we describe our experiences using 3D cell culture to try and assay T cell-mediated cytotoxicity of cancer cell spheroids. We have found that quantifying T cell-mediated cytotoxicity in 3D culture is not simple. Our approach was to use widefield fluorescence microscopy to increase sample size by measuring many spheroids at once. Confocal microscopy has been typically used for imaging similar samples, which greatly limits the number of spheroids that can be assayed. The methods we describe were reliable for assaying drug-mediated cytotoxicity in spheroids, but did not translate consistently to measuring T cell-mediated cytotoxicity.

There are strengths to this system, especially if the goal is to assess the effect of drugs on tumor cells. Small volumes of sample are required (only 10 ul per channel, 30 ul per chip of spheroid suspension or T cell suspension). The small amount of spheroid suspension that is required could be a particular advantage if studying patient-derived organoids, where tumor sample is limited. Adding treatments to the spheroids in the chips is easy and also doesn’t require large volumes. Staining the samples for fluorescent microscopy is relatively easy and short in that no fixation or further manipulation of the samples is required.

Limitations and complications to this system are numerous (listed in Figure 7), especially in terms of T cell-mediated cytotoxicity using exogenous T cells. It is hard to be consistent with the number of spheroids and T cells that are fed into the system. The concentration of T cells can be determined before addition to the chip, but we don’t know how many T cells actually make it into the gel and come in contact with spheroids. Cell death may be overestimated because any dead T cells are also stained with the dead cell stain. In the 2D culture assays we were able to partially control for signal coming from dead T cells by plating wells with T cells alone. Such a strategy would not be possible in 3D culture. However, even in our 2D co-culture assays we may be overestimating cell death if T cells that are in co-culture die more than T cells not in co-culture. If spheroids are the required material, it should be noted that not all cell lines form spheroids equally (some may take longer than others, some may not form spheroids at all). While the treatment and staining protocol is simple, it is hard to estimate the working concentration for many of the reagents used (such as drug treatments, stains, or antibodies). Treatments or stains are pipetted into the side channel/s and diffuse across the central gel channel (becoming diluted). The 3D cell culture chips become expensive if many conditions and replicates are required. Imaging requires a sophisticated fluorescence microscope with automation. Automated quantification of cytotoxicity using fluorescence staining is possible, but not an easy task, and may require model retraining for different types of samples. Lastly, the experiments generate large amounts of data that need to be stored and processed.

**Figure 7:**
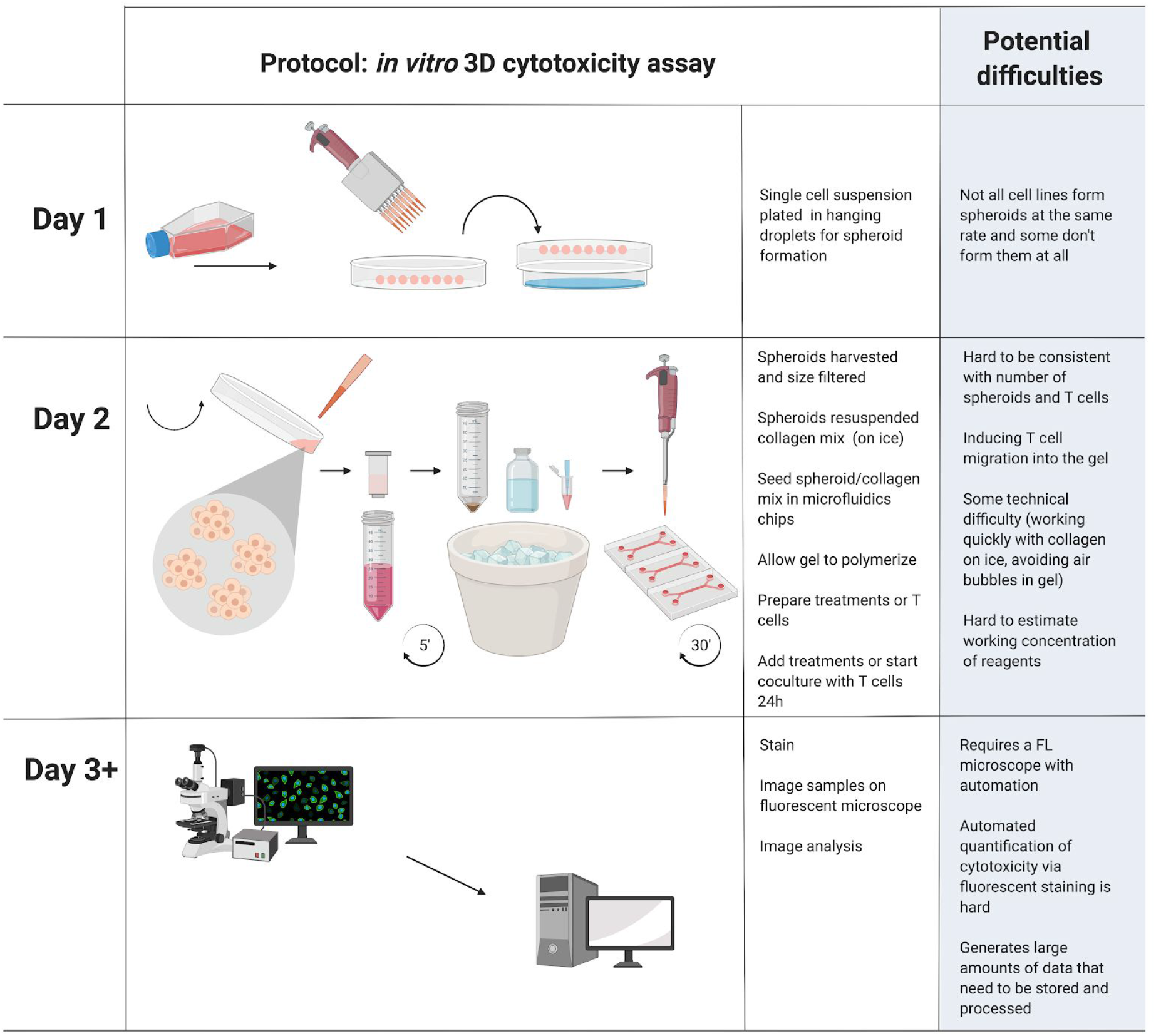
Overview of 3D cytotoxicity assay protocol and points at which potential difficulties can occur.

We had some assumptions about this system that did not appear to be true or that we were unable to address. We had thought that perhaps each spheroid in a gel channel could be thought of as a different sample (meaning we could measure about 300 “tumors” at a time per chip). But through the testing of different computational methods to measure cytotoxicity in the spheroids (in Supplemental Figure 13 and 14), it became apparent that there was no difference between measuring death in the single cells versus in spheroids. The lack of variability between spheroids indicated that maybe we can’t consider each spheroid as a separate tumor sample. However, this could be a result of using a cancer cell line to form the spheroids. Perhaps this would change if patient or mouse-derived tumor material was used to form organoids that retain the heterogeneity of the tumor. Additionally, one of our goals with this system was so have an intermediary assay between 2D cell culture and the mouse model. We had thought cytotoxicity measurements of spheroids in 3D culture could somehow be analogous to measuring the shrinkage of mouse tumors via calipers. But we did not get to a point in our studies where we could make this comparison. We had hoped to be able to measure infiltration properties of T cells with this system, but it looks like it would be hard to differentiate T cells from tumor cells because of the diffraction associated with widefield microscopy. Confocal may be better suited to T cell infiltration questions.

Potential ways this system could be expanded would be to introduce other cells types or ECM components into the tumor microenvironment, as a bottom-up approach. A top-down approach, using mouse- or patient-derived organoids, could be to observe the effect of the tumor microenvironment from a mouse tumor on newly infiltrating T cells. Aside from T cell-tumor interactions, characteristics of T cell migration in 3D culture conditions can be assayed in the absence of tumor cells.

Whether to choose a 2D or 3D cell culture cytotoxicity assay depends on what the research question is. Based our experiences described here, *in vitro* measurement of T cell-mediated cytotoxicity in 2D cell culture produces more reliable and consistent results than in 3D culture, and is a simpler assay to conduct. However, if the research question includes aspects of T cell behaviour such as migration, a 3D cell culture system may be worth trying.

## Materials and Methods

### Cell culture

#### Cancer cell lines

B16F-10 mouse melanoma cells were purchased from ATCC (<u>CRL-6475</u>, lots 70007207 and 70017905). MC38 mouse colon adenocarcinoma cells were kindly provided by the Rubinstein Lab, Medical University of South Carolina. B16-F10 and MC38 cells were cultured in DMEM (HyClone, GE Healthcare, Chicago IL) with 10% fetal bovine serum (Atlas Biologicals, Fort Collins, CO), and 1% penicillin-streptomycin (Thermo Fisher Scientific, Waltham, MA) at 37°C and 5% CO_2_. Cultures were maintained sub-confluency, splitting twice per week with 0.05% trypsin (HyClone).

Culture media protocol: Elinor Gottschalk, Bulent Arman Aksoy, Pinar Aksoy, Jeff Hammerbacher. MC38 and B16-F10 culture media. protocols.io dx.doi.org/10.17504/protocols.io.2y2gfye

#### Mouse primary T cells

Mouse spleens from pmel-1 TCR-transgenic mice were kindly provided by the Paulos lab, Medical University of South Carolina. Isolated splenocytes were frozen at 20 millions cells/ml in Recovery™ Cell Culture Freezing Medium (Thermo Fisher Scientific). When cells were needed for experiments, splenocytes were thawed and cultured in T cell culture media (dx.doi.org/10.17504/protocols.io.qu5dwy6): RPMI with L-glutamine (Corning, Corning, NY), 10% fetal bovine serum (Atlas Biologicals), 715 μM 2-mercaptoethanol (EMD Millipore, Burlington, MA), 25 mM HEPES (HyClone), 1% Penicillin-Streptomycin (Thermo Fisher), 1X sodium pyruvate (HyClone), and 1X non-essential amino acids (HyClone), and supplemented with 200 IU/ml of IL-2 (NCI preclinical repository). On the day that cells were thawed for experiments, hgp100 (GenScript, Piscataway, NJ) was added to the cultures to activate the T cells at a final concentration of 1 μM. Starting at day 3 post-activation, the cultures were counted and expanded if necessary, supplementing with fresh T cell media and 200 IU/ml IL-2 (based on the total volume of the culture) to keep the cells growing at a concentration of 500,000-1 million cells per ml. Cultures were started in 6 well plates or T75 flasks, depending on the starting number of cells, and expanded to T150 flasks.

Protocol details: Bulent Arman Aksoy, Pınar Aksoy, Megan Wyatt, Chrystal M. Paulos, Jeff Hammerbacher (2018). Human primary T cell culture media. protocols.io dx.doi.org/10.17504/protocols.io.qu5dwy6

#### Human primary T cells

Healthy donor blood was purchased from Plasma Consultants LLC (Monroe Township, NJ) from which T cells were isolated with the EasySep™ Direct Human T Cell Isolation Kit (StemCell, Vancouver, Canada). T cells were frozen at 20 million cells/ml in Recovery™ Cell Culture Freezing Medium (Thermo Fisher Scientific).

### 2D T cell-mediated cytotoxicity assay

The day before co-culture with T cells, MC38 or B16-F10 cells were plated at 25,000 cells per well in clear-bottom tissue culture 96 well plates in complete DMEM and incubated overnight (at 37°C and 5% CO_2_). The next day, DMEM was aspirated and pmel-1 T cells that had been cultured for 5 - 11 days were added in T cell media, at the described ratios, to the cancer cells. In the case of MC38 cells, 1 μM of hgp100 peptide (GenScript) in complete DMEM was added for 1 hour at 37°C before co-culture with the pmel-1 T cells. The peptide was washed out with T cell media before T cells were added to the cultures. The co-cultures were incubated overnight. The next day, the plates were assayed using the CytoTox-ONE Homogenous Membrane Integrity Assay (#G7890, Promega, Madison, WI) as per the kit protocol. Briefly, plates were removed from the incubator, lysis buffer was added to the MC38s in the positive control wells, and the plates were allowed to equilibrate to room temperature for 30 minutes. Assay buffer and substrate mix was added to wells and incubated at room temperature for 10 minutes, protected from light. Stop solution was added to wells in the same order and fluorescence was detected with the Spectramax i3x (Molecular Devices LLC, San Jose, CA) with excitation 560 nm and emission 590 nm. Percent cytotoxicity was calculated with the following formula:

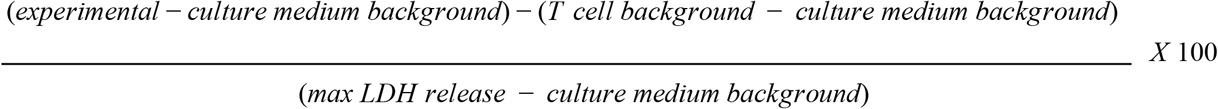

where experimental is the co-culture well, culture medium background is culture media with no cells, T cell background is T cells only at the different concentrations corresponding to the co-culture ratios, and max LDH release is MC38 cells lysed with a detergent to release all of the LDH in the cells into the culture media.

Protocol details: Elinor Gottschalk, Bulent Arman Aksoy, Pinar Aksoy, Jeff Hammerbacher. In vitro pmel-1 T cell-mediated cytotoxicity assay with CytoTox-ONE Homogeneous Membrane Integrity Assay (Promega). protocols.io dx.doi.org/10.17504/protocols.io.6j4hcqw

### Plate-based viability assay

25,000 MC38 cells were plated per well in a 96 well plate and allowed to adhere to the bottom of the wells overnight. The next day, the culture media was exchanged for media containing the antibiotic Staurosporine (MP Biomedicals, LLC, Irvine CA). Staurosporine was diluted in complete DMEM and serially diluted in the wells in the 96 well plate using a multichannel pipette. MC38 cells were incubated overnight with the antibiotic before adding Alamar Blue (Resazurin) (Sigma-Aldrich, St. Louis, MO) solution to the wells. After the addition of Alamar Blue solution, the plates were incubated for another 24 hours. The absorbance at 570 nm and 505 nm were read on a plate reader. The 595 nm reading was subtracted from the 570 nm reading for each well.

Protocol details: Bulent Arman Aksoy, Pınar Aksoy, Jeff Hammerbacher (2018). Resazurin viability assay for human primary T cells in 96-well format. protocols.io dx.doi.org/10.17504/protocols.io.quwdwxe

### 3D culture cytotoxicity assay

A suspension of MC38 cells at 2.5 × 10^6^ cells per ml (in DMEM, 10% FBS, and 1% pen/strep) was pipetted in 20 ul droplets on the surface of a 10 cm tissue culture dish lid using a multichannel pipettor. The lid with the droplets was inverted over the bottom half of the dish, containing 5-10 ml of sterile water to humidify the chamber and incubated overnight 37°C (5% CO_2_). The following day the droplets containing MC38 spheroids were harvested in DMEM and sequentially size filtered through 40 μm and then 100 μm mesh filters (as described by (Jenkins et al. 2018). The selected spheroids between 40 and 100 μm were centrifuged at 200 x g for 5 minutes. A 2.5 mg/ml collagen type I (rat tail, Corning Life Sciences, Corning, NY) gel mixture was prepared using AIM Biotech’s general protocol for gel preparation: https://www.aimbiotech.com/general-protocols.html. IL-2 (at a final concentration of 200 IU per ml and murine CXCL11 (Peprotech, Rocky Hill, NJ) at a final concentration of 2.7 mg/ml were added to the collagen mixture; water volume was adjusted to compensate for these additions in order to maintain the correct collagen concentration). The MC38 spheroid pellet was resuspended in the collagen gel and pipetted into the central channel of 3D cell culture chips (AIM Biotech, Singapore). Sterile water was added to the chambers in the chip holders and the chips were incubated for 30 minutes at 37°C. While the chips incubated, pmel-1 T cells were counted (for these experiments the T cells were 5 - 7 days post activation with the hgp100 peptide). The T cells were centrifuged (350 x g for 5 minutes) and resuspended in T cell culture media (described above) supplemented with IL-2 (200 IU/ml) to achieve the desired concentration of cells per ml. The cells were serially diluted for lower concentrations used in the experiments. Once the gel had polymerized in the microfluidics chips, T cell culture media supplemented with IL-2 (200 IU/ml) and with or without hgp100 (2 μM) was pipetted into the right side channel. T cell suspension was pipetted into the left side channel. In conditions where the spheroids were being killed by Staurosporine (MP Biomedicals), the drug was diluted in complete DMEM and pipetted into both side channels in the chips. The chips were returned to the incubator overnight. After overnight co-culture with T cells or with Staurosporine, the chips were stained for 30 minutes at room temperature in a staining solution containing PBS, 2X NucBlue™ Live ReadyProbes™ Reagent (to stain all nuclei) and SYTOX™ Red Dead Cell Stain (to stain only dead cells) (catalog numbers R37605 and S34859, Thermo Fisher Scientific). The chips were imaged on a Keyence BZ-X710 system (Keyence Corporation, Osaka, Japan) with a 20X objective, DAPI and Cy5 filters and brightfield, and a 20 μm step-size.

Three 3 × 11 grids were used to capture the central channels in a chip. One chip was imaged at a time to reduce the amount of time a chip was exposed to light on the microscope.

Protocol details: Elinor Gottschalk, Bulent Arman Aksoy, Pinar Aksoy, Eric Czech, Jeff Hammerbacher. Image-based 3D cell culture cytotoxicity assay. protocols.io https://protocols.io/view/image-based-3d-cell-culture-cytotoxicity-assay-72shqee

### 3D culture image analysis

Image quantification in this study was conducted using one of three methods and each was applied to the results from all experimental conditions. The first of these identifies individual spheroid objects and integrates all fluorescent signals within those objects. Where not stated explicitly, this method is used for all distribution summaries and statistical comparisons. The two other methods employed extend this process by attempting to further identify individual cells within spheroids. The first of these uses the U-Net-based nuclei segmentation model of CellProfiler (Caicedo et al. 2019) via Cytokit (Czech et al. 2018) while the second utilizes a custom cytometry function based on differences of gaussian filters. This second method affords greater flexibility in parameterization to aid in tuning thresholds (albeit subjectively) to better capture nuclei images with noisy, occluded boundaries. These processes are summarized in **Supplemental Figure 10**.

Multiple methods were applied across culture conditions to ensure that results were less sensitive to computational choices, as well as to provide evidence on the viability of cell quantification within spheroids given that cell images are known to lack penetrance into spheroid cores (Boutin et al. 2018), (Hari et al. 2019). Despite this limitation, we hypothesized that chemically or immunologically mediated cell death would be detectable along spheroid surfaces and that the extent to which it is detectable would correlate with the severity of the mediation. **Supplemental Figure 13** and **Supplemental Figure 14** demonstrate the lack of sensitivity to choice in computational method for cytotoxicity calculations. The first of the three methods above, which does not quantify individual cells, was used for most toxicity measurements reported due to its simplicity and efficiency relative to the other methods.

A component common to all of the methods was the estimation of spheroid object boundaries. Ilastik pixel classification was used to generate probability images before passing them through thresholding and peak local distance functions in order to provide seed points for spheroid centers. Spheroid objects were then separated from one another, when in contact, using the voronoi-based segmentation algorithm of (Jones, Carpenter, and Golland 2005). Pixel classification training was done based on a random sampling of approximately 5 images from each experimental condition (i.e. images at a particular concentration in a titration). This resulted in ~50 images to annotate for each of two separate models trained, one to recognize spheroids in experiments using chemical cytotoxicity (staurosporine) and another to recognize spheroids in T cell co-culture assays.

For individual cell segmentation, the method available via CellProfiler is presented in detail elsewhere (McQuin et al. 2018). The second method however, employing Difference of Gaussian (DoG) kernels, was more onerous in its application and required more manual tuning. Like applications of Laplacian of Gaussian (LoG) filters for cell detection ((Stegmaier et al. 2014), (Al-Kofahi et al. 2010), (Byun et al. 2006)), our method uses DoG as an approximation to the LoG operator in order to improve computational efficiency. This efficiency was important for optimizing the standard deviation range used in the DoG operator, particularly since the operator is applied at multiple scales and individual cells were chosen based on the locations of local maxima created by the "scale space" resulting from the application of the DoG operator at multiple scales. This method is commonly referred to as blob detection (Lindeberg 1993). Cells identified using this method were assumed to match the shape and size implied by the kernel used to identify them.

### T cell migration assay

Collagen gel mixture was prepared as described above and pipetted directly into the central channel of 3D Cell Culture Chips (AIM Biotech). Reconstituted CXCL11 (Peprotech) was added to the gel mixture at the described concentrations. T cells were stained with 0.11 uM IncuCyte^®^ CytoLight Rapid Green Reagent (Essen Bioscience, Ann Arbor, MI) for live cell cytoplasmic labeling for 20 minutes at 37°C. Once the collagen gel had polymerized, T cells were added at a concentration of 10 million cells per ml to the left side channel in the 3D Cell Culture Chips and T cell media supplemented with IL-2 (200 IU/ml) was added to the right side channel. The chips were incubated overnight at 37°C (5% CO_2_). Chips were imaged the following day on the Keyence BZ-X710 system with a 20X objective, GFP filter and brightfield, and a 20 μm step-size. Three 3 × 11 grids were used to capture the central channels in a chip. One chip was imaged at a time to reduce the amount of time a chip was exposed to light on the microscope.

Protocol details: Elinor Gottschalk, Bulent Arman Aksoy, Pinar Aksoy, Jeff Hammerbacher. Staining cells with IncuCyte Cytolight Rapid Dyes for flow cytometry or fluorescent microscopy. protocols.io dx.doi.org/10.17504/protocols.io.73nhqme

### Detecting SIINFEKL presentation

To test the detection of SIINFEKL presentation via fluorescence microscopy in 2D culture, MC38s that had been plated one day prior in 35 mm tissue culture imaging dishes (ibidi, Gräfelfing, Germany) were pulsed with the SIINFEKL peptide at 10 uM for 2 hours at 37°C. The cultures were washed once with PBS and stained. The stain solution contained a 1:200 dilution of an antibody against H-2Kb bound to SIINFEKL conjugated to PE (#141603, Biolegend, San Diego, CA) to detect SIINFEKL peptide being presented at the cell surface, Hoechst to stain the cell nuclei, diluted in PBS with 20% FBS (Atlas Biologicals). The cells were stained for 30 minutes at 37°C before washing twice with PBS. The cultures were imaged in PBS.

To test the duration of peptide presentation in MC38 spheroids in 3D culture, the SIINFEKL peptide at 10 uM was added either to the side channels of 3D cell culture chips (AIM Biotech) with MC38 spheroids in collagen gel in the center, or added directly to the collagen gel/MC38 spheroid mixture before polymerization in the chip. The spheroids were exposed to the peptide for 2 hours or 24 hours. After incubation with the peptide, excess media was removed from the troughs of the side channel ports and 20 ul of staining buffer described above (anti-H-2Kb bound to SIINFEKL and Hoechst in PBS with 20% FBS) was pipetted into the side channels. The spheroids were stained for 30 minutes at 37°C before washing twice with PBS. The cultures were imaged in PBS.

The 2D and 3D cultures were imaged on the Keyence BZ-X710 system with a 20X objective and TRITC filter.

### Flow cytometry

Flow cytometry was performed on the BD FACSVerse (BD Biosciences, San Jose, CA). On the day of analysis, pmel-1 T cells were counted and centrifuged at 350 x g for 5 minutes. The cells were resuspended in PBS with 20% FBS containing the indicated antibodies (listed below) at the concentration recommended by the manufacturer (Biolegend). Cells were stained at room temperature for 20 to 30 minutes, protected from light, and rotating. After staining, the samples were centrifuged, stain buffer removed, and the pellet was resuspended in PBS. Flow cytometry results were analyzed with FlowJo software v10 (Ashland, OR).

### Antibodies

| Name | Vendor | Catalog # |
| --- | --- | --- |
| CD44 | Biolegend | 103040 |
| CD62L | Biolegend | 104412 |
| CD69 | Biolegend | 104532 |
| CD25 | Biolegend | 101918 |

## Acknowledgments

We thank the Paulos Lab and the Rubinstein Lab at the Medical University of South Carolina for providing us with pmel-1 splenocytes and MC38 cells, respectively. This work made use of the Flow Cytometry and Cell Sorting Unit Shared Resource, Hollings Cancer Center, Medical University of South Carolina (P30 CA138313). The authors declare no competing financial interests.

## Data availability

Intermediate and final data sets that were used to generate the figures and summaries in this manuscript are available at https://github.com/hammerlab/spheroid. All raw spheroid image data sets are publicly available under gs://cytokit/datasets/spheroids.

## References

1. Abad, John D., Claudia Wrzensinski, Willem Overwijk, Moniek A. De Witte, Annelies Jorritsma, Cary Hsu, Luca Gattinoni, et al. 2008. “T-Cell Receptor Gene Therapy of Established Tumors in a Murine Melanoma Model.” Journal of Immunotherapy 31 (1): 1–6.

2. Abu-Shah, Enas, Philippos Demetriou, Viveka Mayya, Stefan Balint, Mikhail A. Kutuzov, Omer Dushek, and Michael L. Dustin. 2019. “A Tissue-like Platform for Studying Engineered Quiescent Human T-Cells’ Interactions with Dendritic Cells.” bioRxiv. https://doi.org/10.1101/587386.

3. Aksoy, Pinar, Bülent Arman Aksoy, Eric Czech, and Jeff Hammerbacher. 2019. “Viable and Efficient Electroporation-Based Genetic Manipulation of Unstimulated Human T Cells.” bioRxiv. https://doi.org/10.1101/466243.

4. Al-Kofahi, Yousef, Wiem Lassoued, William Lee, and Badrinath Roysam. 2010. “Improved Automatic Detection and Segmentation of Cell Nuclei in Histopathology Images.” IEEE Transactions on Bio-Medical Engineering 57 (4): 841–52.

5. Belmokhtar, C. A., J. Hillion, and E. Ségal-Bendirdjian. 2001. “Staurosporine Induces Apoptosis through Both Caspase-Dependent and Caspase-Independent Mechanisms.” Oncogene 20 (26): 3354–62.

6. Berg, Stuart, Dominik Kutra, Thorben Kroeger, Christoph N. Straehle, Bernhard X. Kausler, Carsten Haubold, Martin Schiegg, et al. 2019. “Ilastik: Interactive Machine Learning for (bio)image Analysis.” Nature Methods, September. https://doi.org/10.1038/s41592-019-0582-9.

7. Boorn, Jasper G. van den, Debby Konijnenberg, Esther P. M. Tjin, Daisy I. Picavet, Nico J. Meeuwenoord, Dmitri V. Filippov, J. P. Wietze van der Veen, Jan D. Bos, Cornelis J. M. Melief, and Rosalie M. Luiten. 2010. “Effective Melanoma Immunotherapy in Mice by the Skin-Depigmenting Agent Monobenzone and the Adjuvants Imiquimod and CpG.” PloS One 5 (5): e10626.

8. Boutin, Molly E., Ty C. Voss, Steven A. Titus, Kennie Cruz-Gutierrez, Sam Michael, and Marc Ferrer. 2018. “A High-Throughput Imaging and Nuclear Segmentation Analysis Protocol for Cleared 3D Culture Models.” Scientific Reports 8. https://doi.org/10.1038/s41598-018-29169-0.

9. Byun, Jiyun, Mark R. Verardo, Baris Sumengen, Geoffrey P. Lewis, B. S. Manjunath, and Steven K. Fisher. 2006. “Automated Tool for the Detection of Cell Nuclei in Digital Microscopic Images: Application to Retinal Images.” Molecular Vision 12 (August): 949–60.

10. Caicedo, Juan C., Jonathan Roth, Allen Goodman, Tim Becker, Kyle W. Karhohs, Matthieu Broisin, Csaba Molnar, et al. 2019. “Evaluation of Deep Learning Strategies for Nucleus Segmentation in Fluorescence Images.” Cytometry. Part A: The Journal of the International Society for Analytical Cytology 95 (9): 952–65.

11. Chung, Soobin, Gippeum J. Lim, and Ji Youn Lee. 2019. “Quantitative Analysis of Melanin Content in a Three-Dimensional Melanoma Cell Culture.” Scientific Reports 9 (1): 780.

12. Contento, Rita Lucia, Barbara Molon, Cedric Boularan, Tullio Pozzan, Santos Manes, Stefano Marullo, and Antonella Viola. 2008. “CXCR4-CCR5: A Couple Modulating T Cell Functions.” Proceedings of the National Academy of Sciences of the United States of America 105 (29): 10101–6.

13. Courau, Tristan, Julie Bonnereau, Justine Chicoteau, Hugo Bottois, Romain Remark, Laura Assante Miranda, Antoine Toubert, et al. 2019. “Cocultures of Human Colorectal Tumor Spheroids with Immune Cells Reveal the Therapeutic Potential of MICA/B and NKG2A Targeting for Cancer Treatment.” Journal for Immunotherapy of Cancer 7 (1): 74.

14. Czech, Eric, Bulent Arman Aksoy, Pinar Aksoy, and Jeffrey Hammerbacher. 2018. “Cytokit: A Single-Cell Analysis Toolkit for High Dimensional Fluorescent Microscopy Imaging.” bioRxiv. https://doi.org/10.1101/460980.

15. Dangles-Marie, Virginie, Sophie Richon, Mohamed El-Behi, Hamid Echchakir, Guillaume Dorothée., Jérôme Thiery, Pierre Validire, et al. 2003. “A Three-Dimensional Tumor Cell Defect in Activating Autologous CTLs Is Associated with Inefficient Antigen Presentation Correlated with Heat Shock Protein-70 down-Regulation.” Cancer Research 63 (13): 3682–87.

16. Deng, Jiehui, Eric S. Wang, Russell W. Jenkins, Shuai Li, Ruben Dries, Kathleen Yates, Sandeep Chhabra, et al. 2018. “CDK4/6 Inhibition Augments Antitumor Immunity by Enhancing T-Cell Activation.” Cancer Discovery 8 (2): 216–33.

17. Edmondson, Rasheena, Jessica Jenkins Broglie, Audrey F. Adcock, and Liju Yang. 2014. “Three-Dimensional Cell Culture Systems and Their Applications in Drug Discovery and Cell-Based Biosensors.” Assay and Drug Development Technologies 12 (4): 207–18.

18. Hari, Neelam, Priyanka Patel, Jacqueline Ross, Kevin Hicks, and Frédérique Vanholsbeeck. 2019. “Optical Coherence Tomography Complements Confocal Microscopy for Investigation of Multicellular Tumour Spheroids.” Scientific Reports 9 (1): 1–11.

19. Ho, William Y., Joseph N. Blattman, Michelle L. Dossett, Cassian Yee, and Philip D. Greenberg. 2003. “Adoptive Immunotherapy: Engineering T Cell Responses as Biologic Weapons for Tumor Mass Destruction.” Cancer Cell 3 (5): 431–37.

20. Hu, Joyce K., Takashi Kagari, Jonathan M. Clingan, and Mehrdad Matloubian. 2011. “Expression of Chemokine Receptor CXCR3 on T Cells Affects the Balance between Effector and Memory CD8 T-Cell Generation.” Proceedings of the National Academy of Sciences of the United States of America 108 (21): E118–27.

21. Jenkins, Russell W., Amir R. Aref, Patrick H. Lizotte, Elena Ivanova, Susanna Stinson, Chensheng W. Zhou, Michaela Bowden, et al. 2018. “Ex Vivo Profiling of PD-1 Blockade Using Organotypic Tumor Spheroids.” Cancer Discovery 8 (2): 196–215.

22. Jones, Thouis R., Anne Carpenter, and Polina Golland. 2005. “Voronoi-Based Segmentation of Cells on Image Manifolds.” Computer Vision for Biomedical Image Applications. https://doi.org/10.1007/11569541_54.

23. June, Carl H. 2007. “Adoptive T Cell Therapy for Cancer in the Clinic.” The Journal of Clinical Investigation 117 (6): 1466–76.

24. Knowles, H. J., and R. M. Phillips. 2001. “Identification of Differentially Expressed Genes in Experimental Models of the Tumor Microenvironment Using Differential Display.” Anticancer Research 21 (4A): 2305–11.

25. Lindeberg, Tony. 1993. “Detecting Salient Blob-like Image Structures and Their Scales with a Scale-Space Primal Sketch: A Method for Focus-of-Attention.” International Journal of Computer Vision. https://doi.org/10.1007/bf01469346.

26. McQuin, Claire, Allen Goodman, Vasiliy Chernyshev, Lee Kamentsky, Beth A. Cimini, Kyle W. Karhohs, Minh Doan, et al. 2018. “CellProfiler 3.0: Next-Generation Image Processing for Biology.” PLoS Biology 16 (7): e2005970.

27. Müller, Ines, and Dagmar Kulms. 2018. “A 3D Organotypic Melanoma Spheroid Skin Model.” Journal of Visualized Experiments: JoVE, no. 135 (May). https://doi.org/10.3791/57500.

28. Overwijk, Willem W., Marc R. Theoret, Steven E. Finkelstein, Deborah R. Surman, Laurina A. de Jong, Florry A. Vyth-Dreese, Trees A. Dellemijn, et al. 2003. “Tumor Regression and Autoimmunity after Reversal of a Functionally Tolerant State of Self-Reactive CD8+ T Cells.” The Journal of Experimental Medicine 198 (4): 569–80.

29. Pavesi, Andrea, Anthony T. Tan, Sarene Koh, Adeline Chia, Marta Colombo, Emanuele Antonecchia, Carlo Miccolis, et al. 2017. “A 3D Microfluidic Model for Preclinical Evaluation of TCR-Engineered T Cells against Solid Tumors.” JCI Insight 2 (12). https://doi.org/10.1172/jci.insight.89762.

30. Qin, S., J. B. Rottman, P. Myers, N. Kassam, M. Weinblatt, M. Loetscher, A. E. Koch, B. Moser, and C. R. Mackay. 1998. “The Chemokine Receptors CXCR3 and CCR5 Mark Subsets of T Cells Associated with Certain Inflammatory Reactions.” The Journal of Clinical Investigation 101 (4): 746–54.

31. Riss, Terry, Andrew Niles, Rich Moravec, Natashia Karassina, and Jolanta Vidugiriene. 2019. “Cytotoxicity Assays: In Vitro Methods to Measure Dead Cells.” In Assay Guidance Manual, edited by G. Sitta Sittampalam, Abigail Grossman, Kyle Brimacombe, Michelle Arkin, Douglas Auld, Christopher P. Austin, Jonathan Baell, et al. Bethesda (MD): Eli Lilly & Company and the National Center for AdvancingTranslational Sciences.

32. Stegmaier, Johannes, Jens C. Otte, Andrei Kobitski, Andreas Bartschat, Ariel Garcia, G. Ulrich Nienhaus, Uwe Strähle, and Ralf Mikut. 2014. “Fast Segmentation of Stained Nuclei in Terabyte-Scale, Time Resolved 3D Microscopy Image Stacks.” PloS One 9 (2): e90036.

33. Theos, Alexander C., Steven T. Truschel, Graça Raposo, and Michael S. Marks. 2005. “The Silver Locus Product Pmel17/gp100/Silv/ME20: Controversial in Name and in Function.” Pigment Cell Research / Sponsored by the European Society for Pigment Cell Research and the International Pigment Cell Society 18 (5): 322–36.

