## Supplemental Figures for "Towards a scaled-up T cell-mediated cytotoxicity assay in 3D cell culture using microscopy"

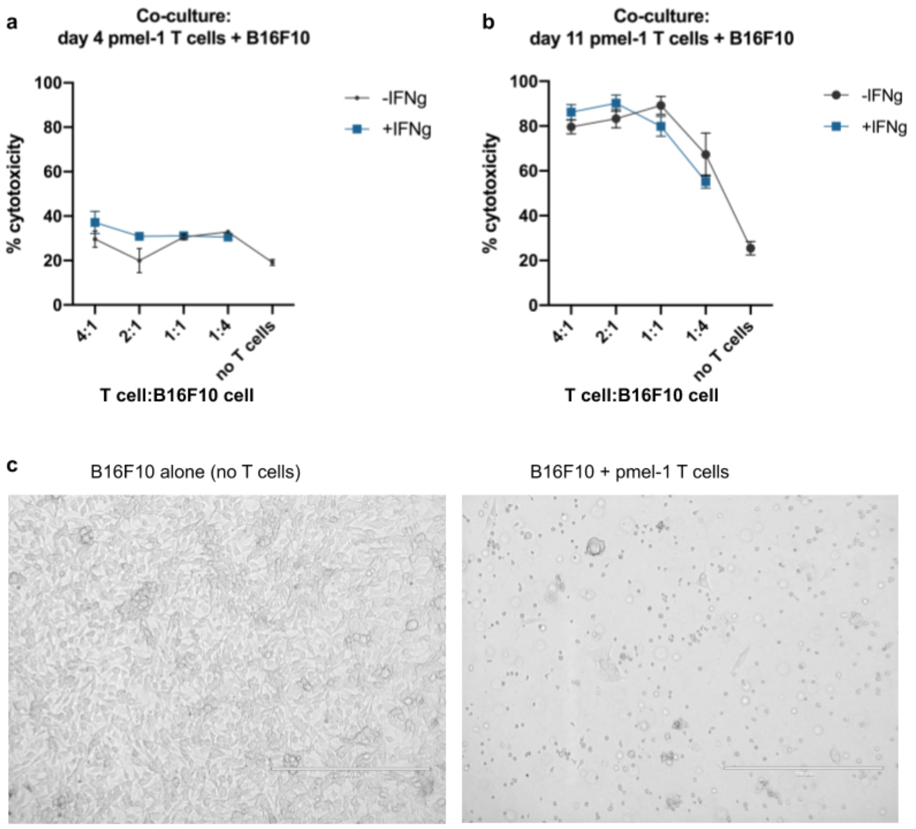


**Supplemental Figure 1: Cytotoxicity assays (LDH release) measuring B16F10 cell death after 2D overnight co-culture with pmel-1 T cells. a)** Day 4 pmel-1 T cells do not kill B16F10 cells, even with IFNγ pre-treatment. **b)** Day 11 Pmel-1 T cells kill B16F10 cells, with or without IFNγ pre-treatment. **c)** Light microscope images of B16F10 cells without pmel-1 T cells (left) and after co-culture with the T cells.

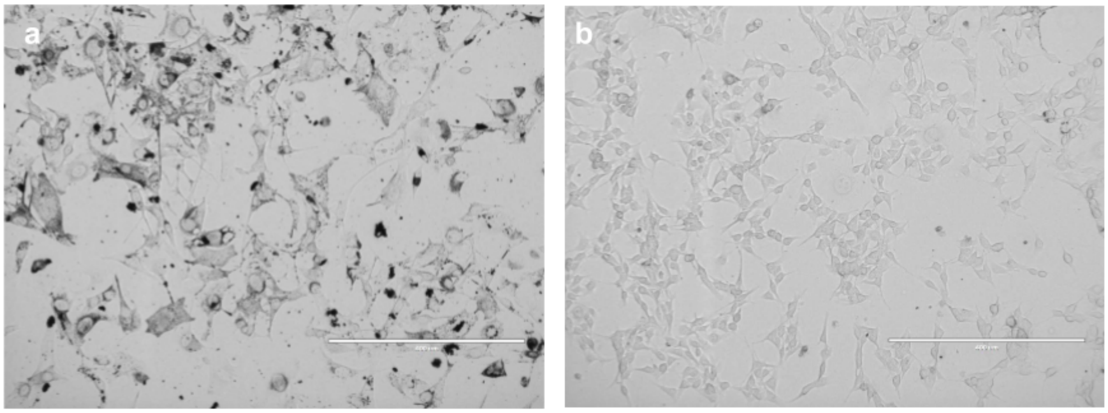


**Supplemental Figure 2: a)** B16F10 cells at passage 2 cultured in DMEM and **b)** at passage 33.

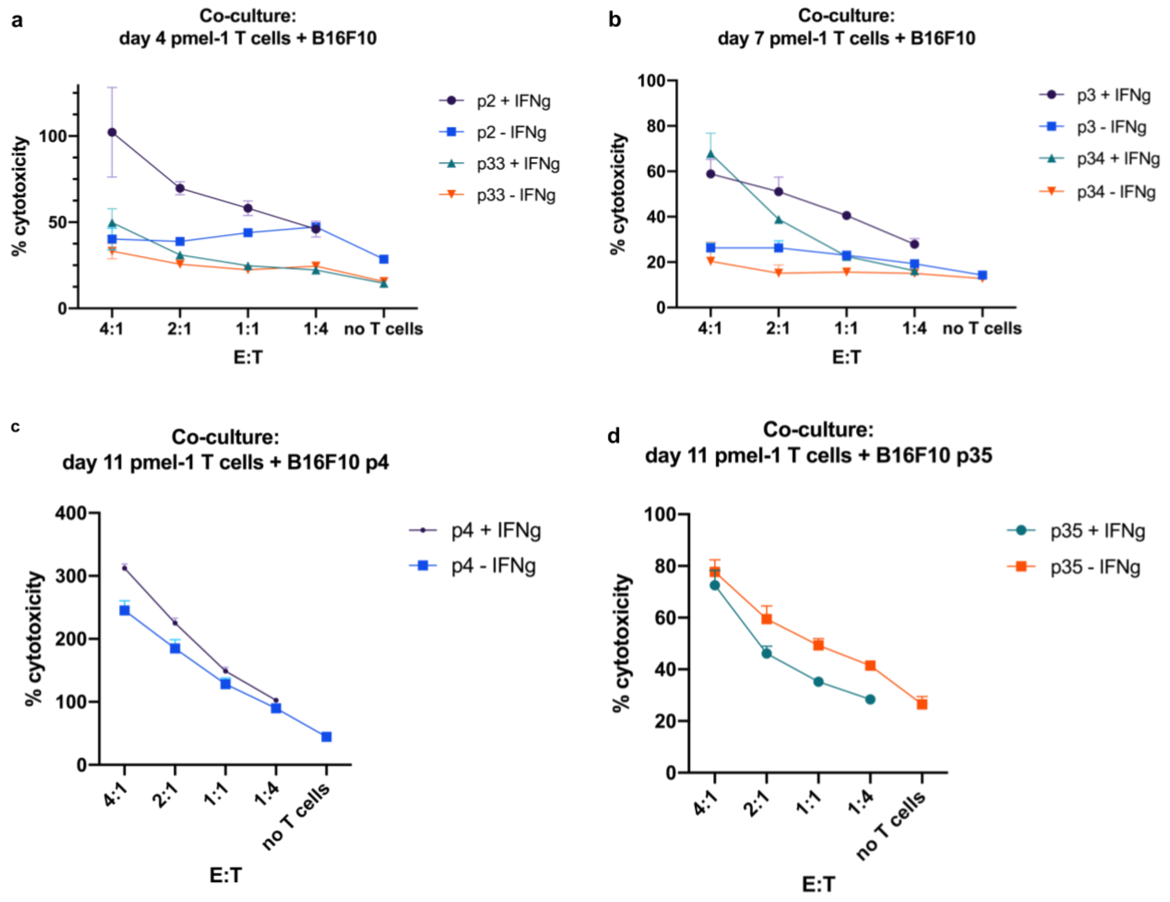


**Supplemental Figure 3: Cytotoxicity assays (LDH release) measuring cell death in early and late passage B16F10 cells after 2D overnight co-culture with pmel-1 T cells. a)** Day 4 pmel-1 T cells only killed early passage B16F10 cells when they had been pre-treated with IFNγ. **b)** Day 7 pmel-1 T cells could kill early and late passage B16F10 cells if they had been pre-treated with IFNγ. **c-d)** Day 11 Pmel-1 T cells kill B16F10, with or without IFNγ pre-treatment.

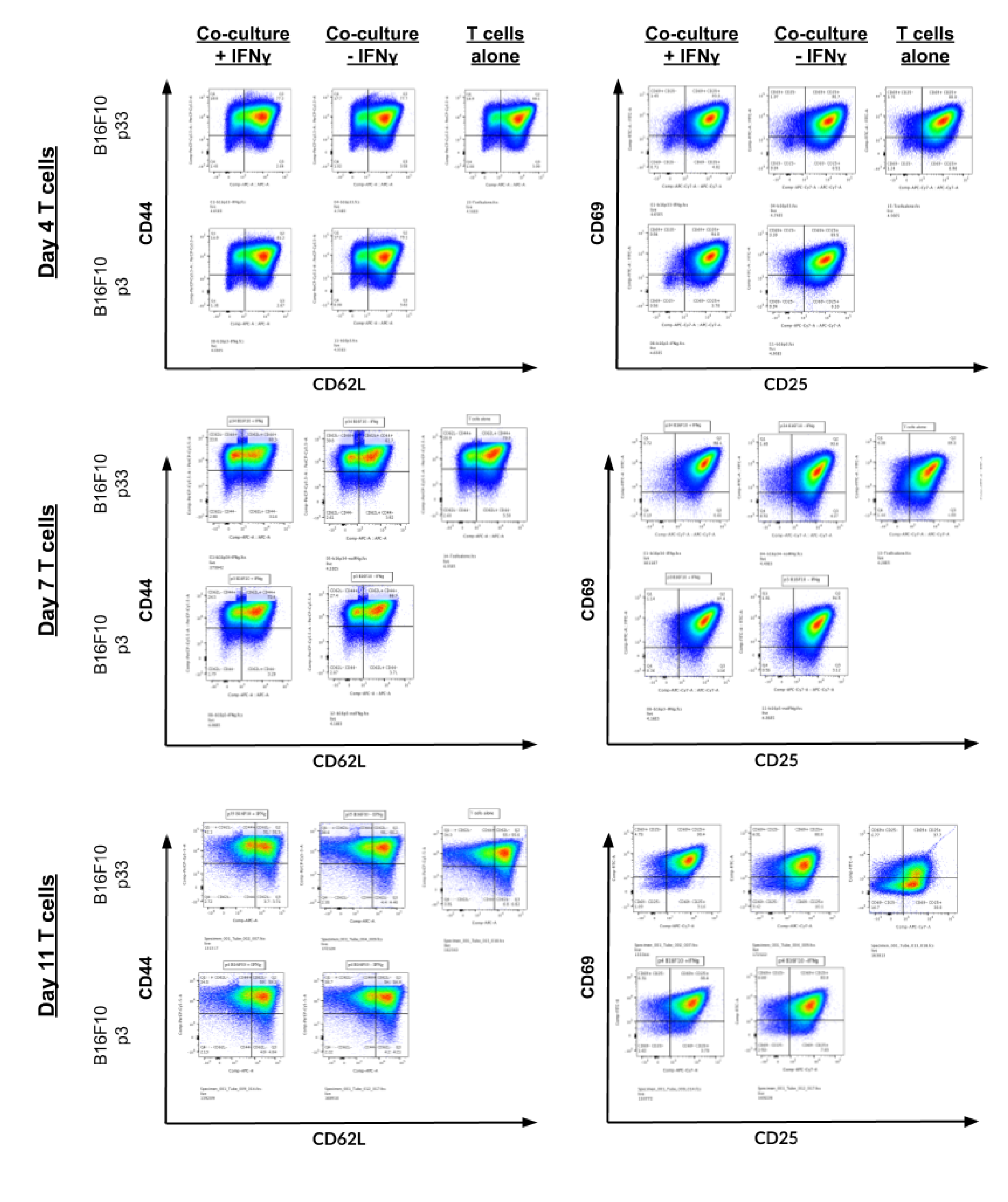

**Supplemental Figure 4: Flow cytometry profiles of pmel-1 T cells (days 4, 7, and 11 post-activation) after overnight co-culture with early and late passage B16F10 cells.**

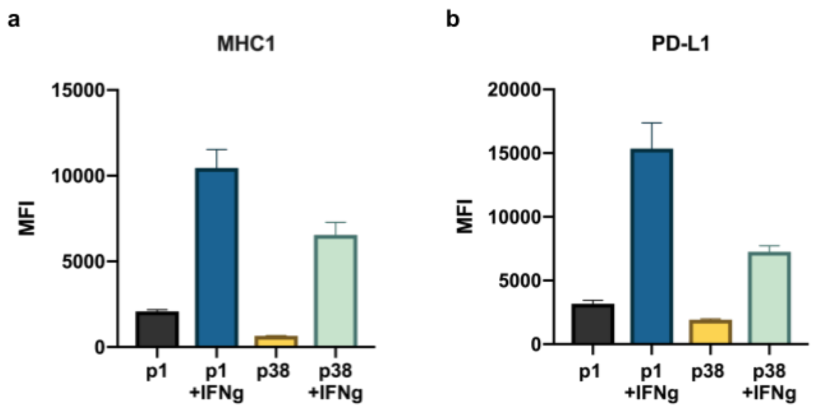


**Supplemental Figure 5: MHCI (a) and PD-L1 (b) levels in early (p1) and late (p38) B16F10 cells with and without overnight IFNγ treatment.** Protein levels were measured via flow cytometry.

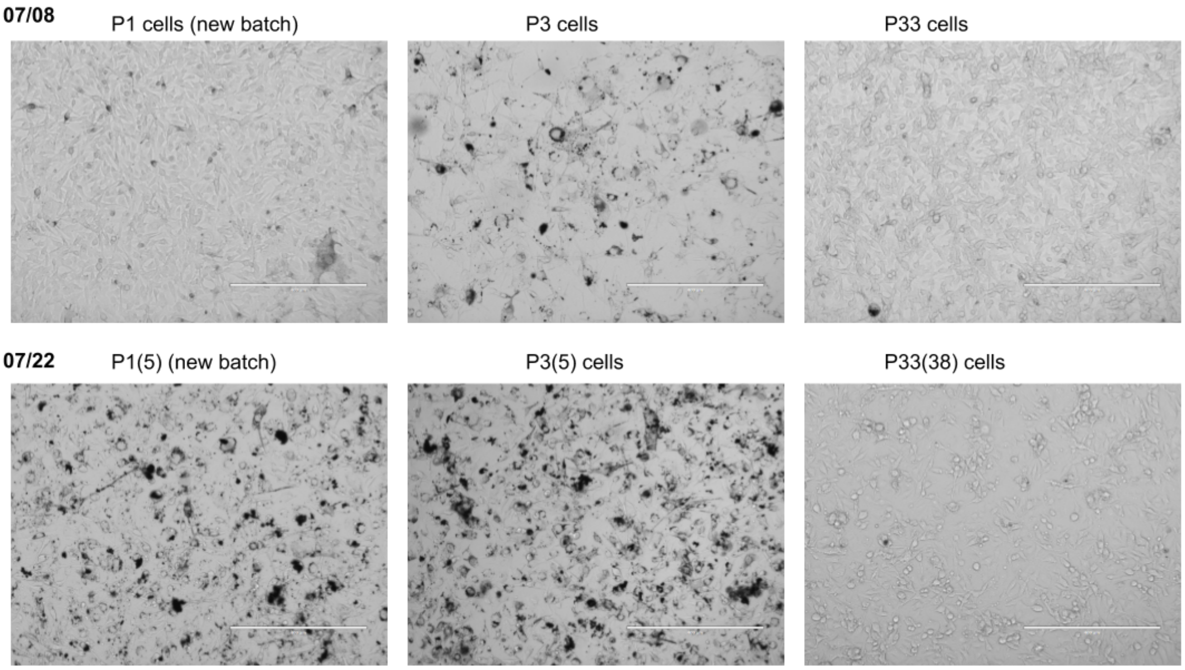


**Supplemental Figure 6: B16F10 cells cultured in DMEM.** Two early passage numbers (different lots from the ATCC: new batch is lot #70017905 and p3 and p33 cells were grown from lot #70007207).

Light microscope images were taken of the cultures on 07/08/2019 and 07/22/2019 at the indicated passage numbers.

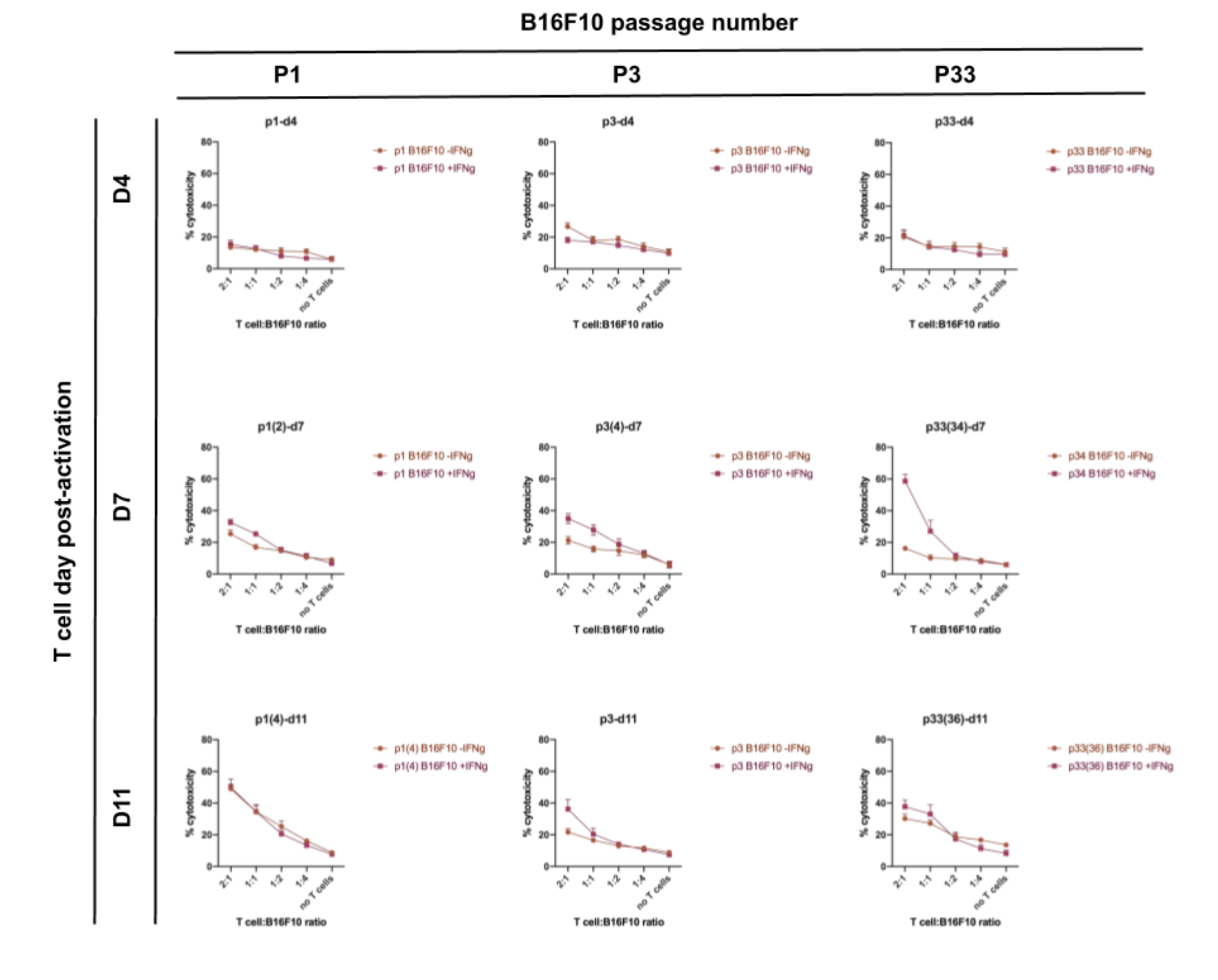


**Supplemental Figure 7: Cytotoxicity assays (LDH release) measuring cell death in early (two different lots) and late passage B16F10 cells after 2D overnight co-culture with pmel-1 T cells.**

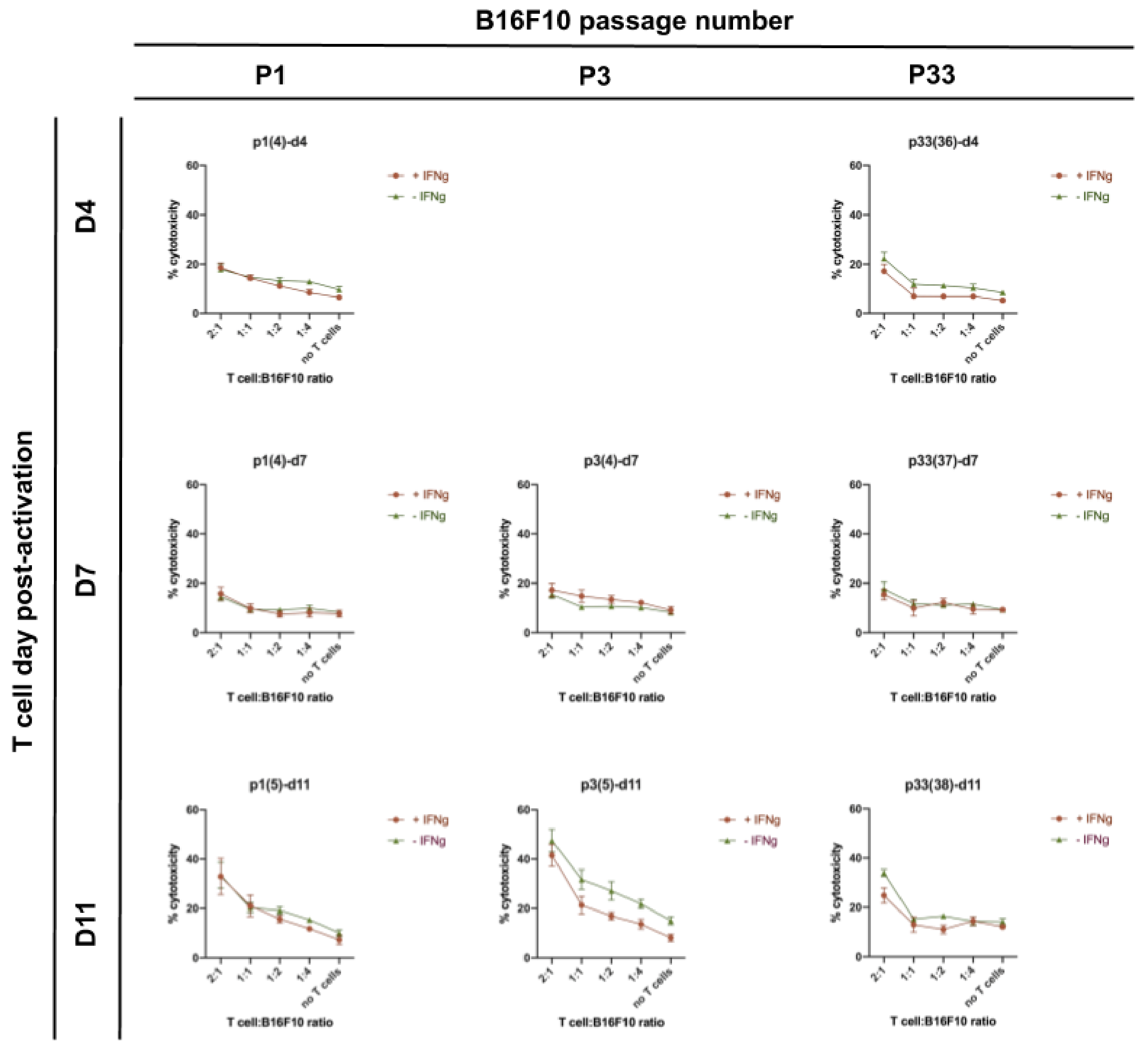


**Supplemental Figure 8: Cytotoxicity assays (LDH release) measuring cell death in early (two different lots) and late passage B16F10 cells after 2D overnight co-culture with pmel-1 T cells.**

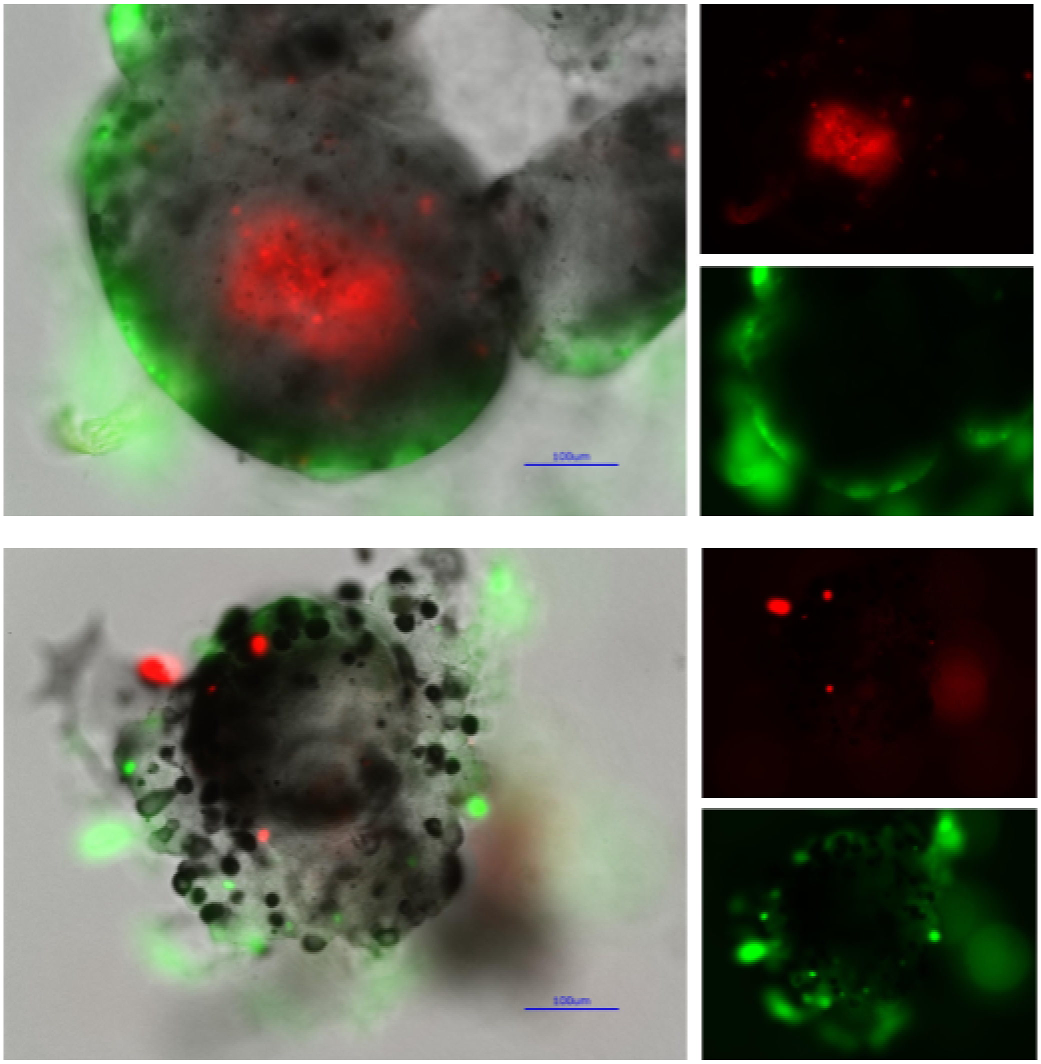

**Supplemental Figure 9:** B16F10 spheroids 22 days after being plated for spheroid formation in hanging droplets and growing for 7 days in collagen gel.

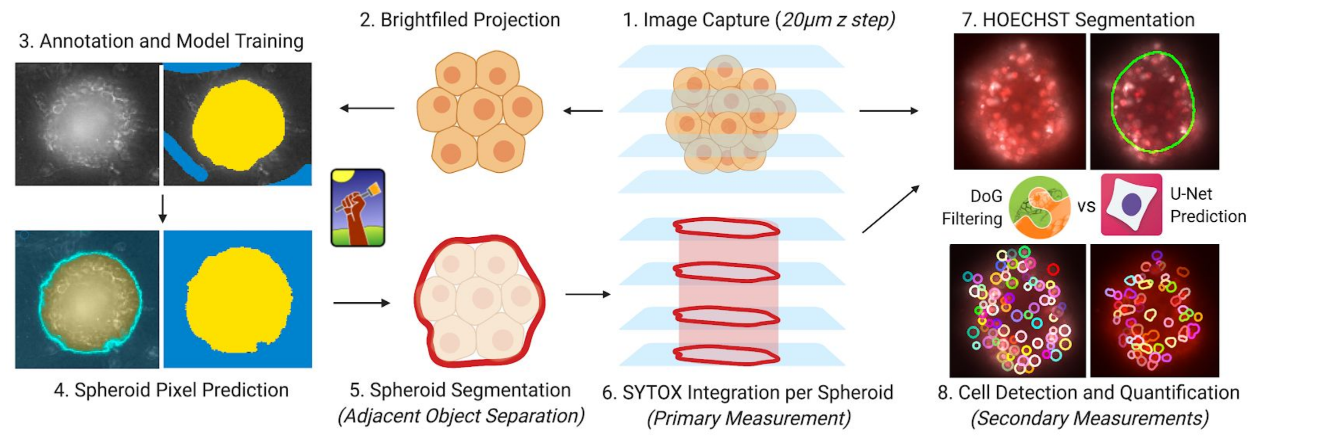


**Supplemental Figure 10: Overview of image quantification process**. Brightfield images are first max-projected to 2D before Ilastik annotation with two-class pixel prediction workflow (spheroid and background). Pixel predictions from trained Ilastik models are applied to produce probability images that are binarized with Hysteresis thresholding (with low threshold .5 and high threshold .8). Spheroid objects are determined based on voronoi-based segmentation over brightfield images using peak local maxima in the distance image corresponding to the threshold image as seed points. The resulting spheroid objects are projected back into 3D, circumscribing an identical 2D area in each z plane, and SYTOX intensities over these 3D volumes are then integrated to produce individual spheroid measurements. Secondary measurements were also collected based on a further segmentation of the spheroid images into individual cells using either a deep learning model provided as part of CellProfiler or a manually parameterized blob detection method. Individual cells propagated to later stages of the analysis were chosen amongst all z planes based on which plane had the most identifiable cells (all others are ignored).

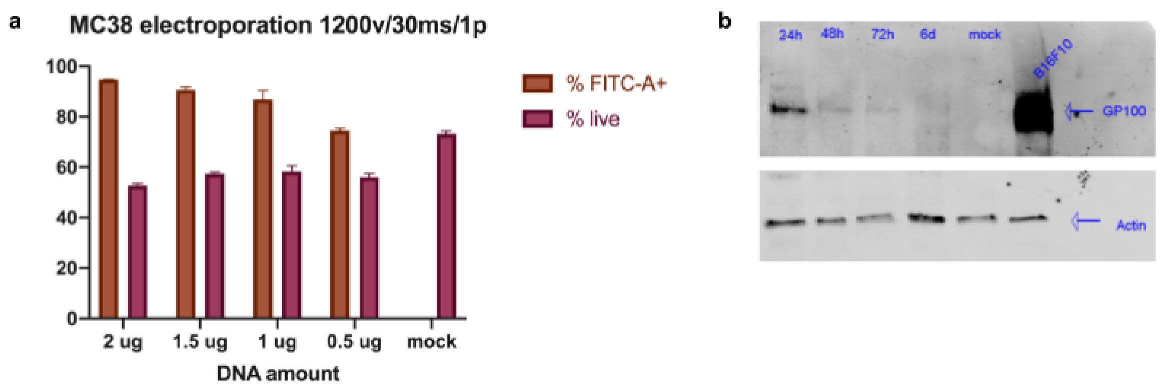


**Supplemental Figure 11: a)** Electroporation efficiency and viability in MC38 cells with different amounts of DNA encoding GFP at the indicated electroporation settings. **b)** Western blot of MC38 lysates at different timepoints post-electroporation with DNA encoding the pmel protein. Blot was probed with anti-GP100 and anti-Actin antibodies.

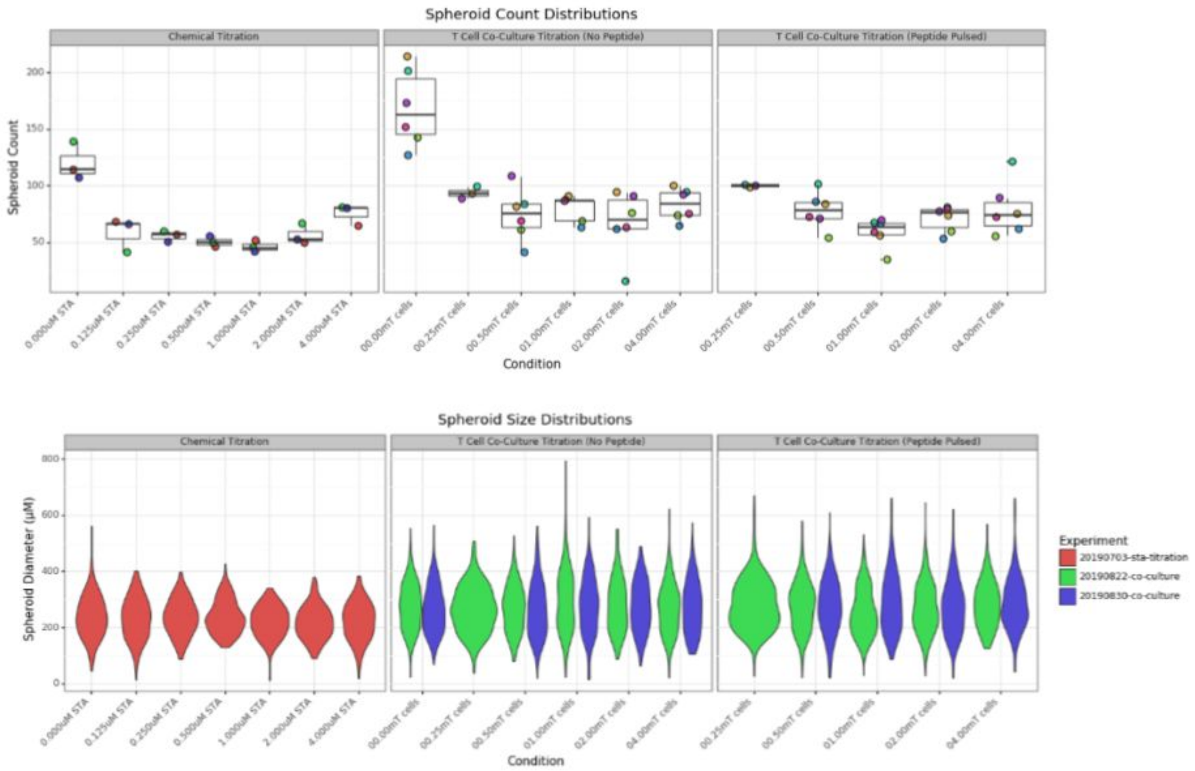


**Supplemental Figure 12: Spheroid size and counts for staurosporine titration and co-culture experiments.** These figures show stability of spheroid counts and sizes across conditions and experiments, with the exception of spheroid counts being larger in untreated groups. The spheroid count distributions are across the individual counts associated with all experiments and replicates that matched the condition (i.e. STA concentration or T cell count).

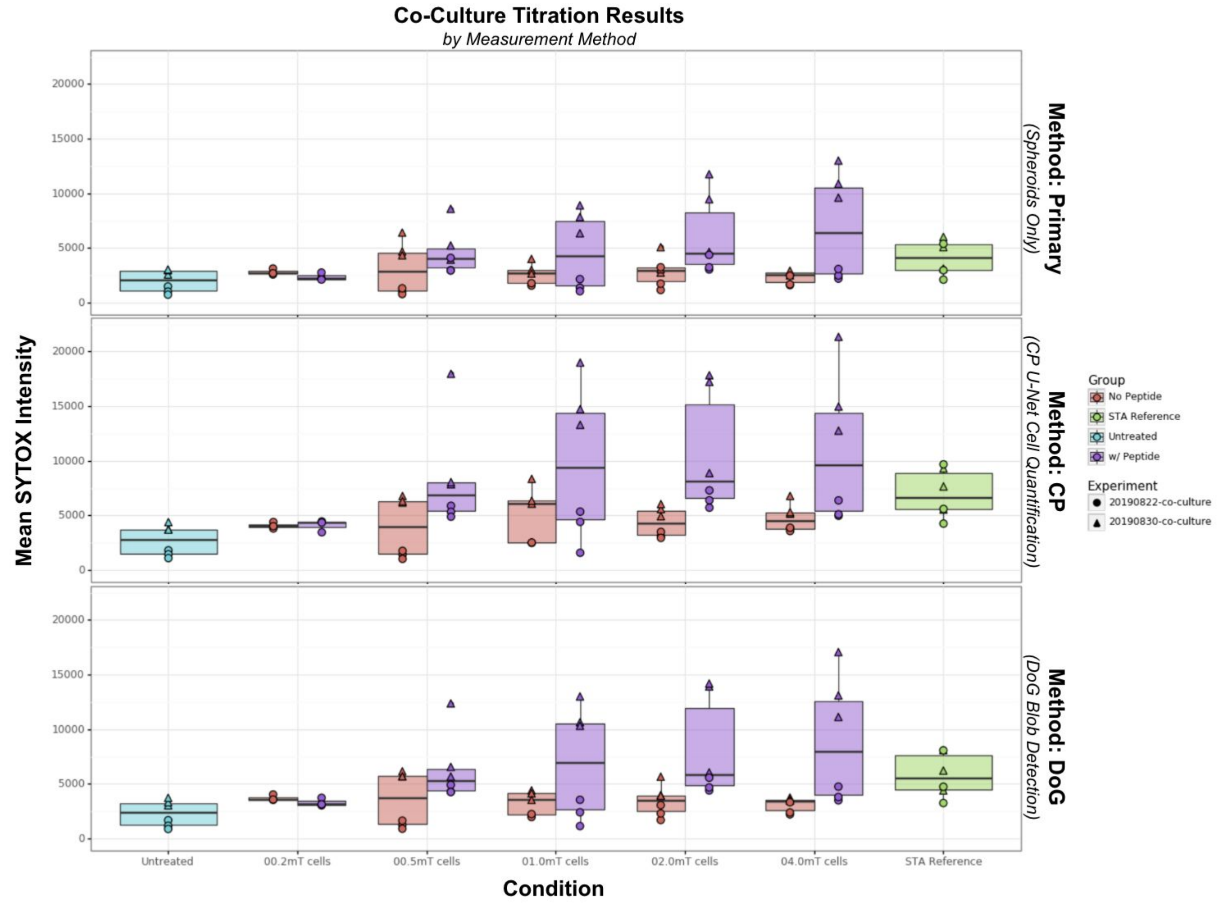


**Supplemental Figure 13: Co-culture titration result invariance to computational method.** These results match those of **Figure 6** (for "Method: Primary") but also show the same results from two extra computational approaches. These methods rely on quantifications of individual cells rather than spheroids, but show little difference in cytotoxicity inferred as a function of the number of T cells introduced into the 3D culture alongside MC38 cells.

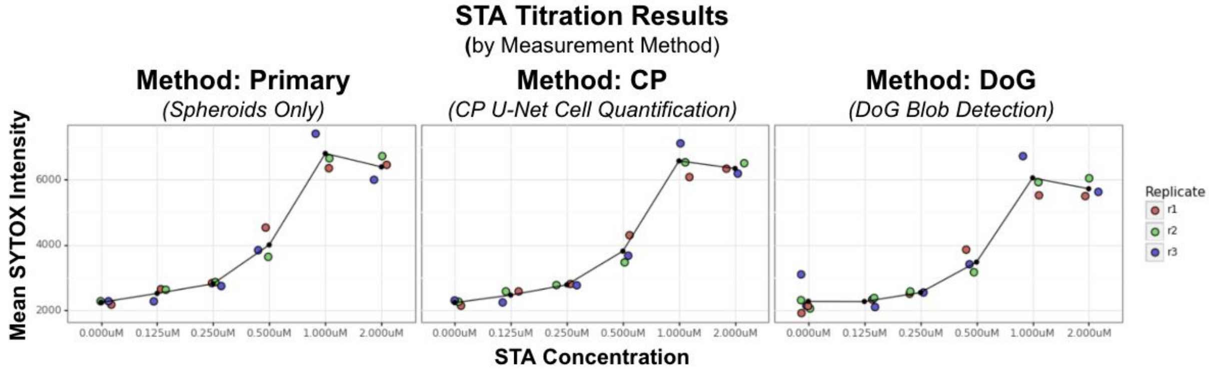


**Supplemental Figure 14: STA titration result invariance to computational method.** These results match those of **Figure 3** (for “Method: Primary”) but also show the same results from two extra computational approaches.  These methods rely on quantifications of individual cells rather than spheroids, but show little difference in cytotoxicity inferred as a function of chemical concentration.
